# Comprehensive Evaluation of AlphaFold-Multimer, AlphaFold3 and ColabFold, and Scoring Functions in Predicting Protein-Peptide Complex Structures

**DOI:** 10.1101/2024.11.11.622992

**Authors:** Negin Manshour, Jarett Zida Ren, Farzaneh Esmaili, Erik Bergstrom, Dong Xu

## Abstract

Determining the three-dimensional structures of protein-peptide complexes is crucial for elucidating biological processes and designing peptide-based drugs. Protein-peptide docking has become essential for predicting complex structures. AlphaFold-Multimer, ColabFold and AlphaFold3 provided groundbreaking tools to enhance the protein-peptide docking accuracy. This study evaluates these three tools for predicting protein-peptide complex structures using Template-Based (TB) and Template-Free (TF) methods. AlphaFold-Multimer excels in TB predictions and performs moderately in TF scenarios in the prediction pool, but TF outperforms TB in the first-ranked models. ColabFold demonstrates versatility in both TB and TF settings. AlphaFold3 generates high-quality structures for more proteins, but the medium accuracy is not as good as AlphaFold-Multimer using a large model pool. We also assessed the performance of various scoring functions in ranking predicted protein-peptide complex structures. While the scoring function built in AlphaFold demonstrates the best performance, some other scoring functions, e.g., FoldX-Stability and HADDOCK-mdscore, provide complementary values. The findings suggest the potential for enhancing scoring functions targeting AlphaFold-based predictions by combining multiple scoring functions or using a consensus approach from many prediction models.

## Introduction

Protein-peptide interactions have been recognized as playing crucial roles in a wide range of biological processes. The participating peptides typically consist of fewer than 50 residues. Approximately 15-40% of protein-protein interaction activities are estimated to involve protein-peptide complexes[1]. Consequently, peptides have increasingly attracted attention as promising binding candidates for drug design, owing to their biochemical properties and relatively low toxicity[2]. Acquiring three-dimensional (3D) structural data of protein complexes is crucial for understanding their functional roles and malfunctions associated with diseases[3–6]. This understanding is essential for elucidating molecular mechanisms underlying protein-peptide recognition and advancing the development of peptide-based therapeutics[7]. To experimentally determine 3D structures of protein-peptide complexes, common techniques include cryo-electron microscopy and X-ray crystallography. While effective, these experimental methods often require extensive labour and costs[8]. Additionally, the experimental characterization of protein-peptide complex structures is considerably hindered by the dynamic and transient nature of some interactions[9]. Due to these challenges, computational methods, such as protein-peptide docking, offer valuable information for protein-peptide complex structures. Docking algorithms fall into two main categories: template-based (TB) and template-free (TF)[1, 10]. In TB docking, known complex structures are used as templates to predict interactions between proteins and peptides, which is effective when similar complex structures are accessible. In contrast, TF docking, also referred to as *ab initio* docking, predicts interactions based solely on the physical and chemical properties of the molecules[11].

Previously, the prediction of protein complex structures was primarily achieved through geometry docking[12]. However, the advent of artificial intelligence (AI), particularly through applications like AlphaFold-Multimer (AFM)[13], has recently revolutionized the field by cofolding protein-protein and protein-peptide complexes[14]. ColabFold (CF) simplifies the use of AlphaFold and AlphaFold-Multimer with Google Colab, and it predicts five TB or TF models for each protein complex. CF greatly improves the speed and ease of predicting protein complexes, thus facilitating fast advancements in structural biology[15]. AlphaFold3 (AF3), the latest version of AlphaFold, is accessible through its dedicated website. It is capable of predicting protein complex structures within five models[16]. Numerous studies have assessed the performance of AlphaFold; however, they concentrated on single protein or protein-protein complexes[17, 18] (including antibody-antigen structures[19, 20]), rather than on protein-peptide complexes. In this study, we focused on assessing the predicted structures of protein-peptide complexes by AFM, CF, and AF3.

Another significant challenge in computational methods is the accurate identification of near-native protein-peptide interaction conformations from a vast array of generated models, a process commonly known as scoring[21]. Several methods have been introduced to rank protein complex structures[22]. The evolution of scoring functions in protein structure prediction has shifted from physics-based methods—exemplified by HADDOCK[23]–and empirical-based methods like AutoDock Vina[24], to knowledge-based potentials, such as FoldX[25]. Physics-based scoring functions began with force-field methods based on fundamental physical principles, incorporating solvent effects and charge features, but these methods are computationally intensive[26]. On the other hand, empirical methods use simplified models of molecular interactions derived from experimental data, resulting in faster computations[27]. Knowledge-based functions use statistical potentials derived from known structural data, balancing accuracy and speed[28].

Scoring functions have further advanced with the adoption of deep learning (DL) strategies, including DOVE[29], GNN_DOVE[30], DeepRank[31], and the graph neural-network-based methods such as DeepRank-GNN[32], DeepRank-GNN-esm[33], and Interpeprank[34]. DL-based scoring functions have been reviewed and assessed in various studies, such as those involving antibody-antigen and enzyme-inhibitor interactions[28]. Other studies have offered a comprehensive comparison of DL-based scoring functions for virtual screening[35] and ranking of the predicted protein-ligand complexes[36]. While numerous scoring functions for protein complex structure assessment have been developed, their performance depends on the methods of predicting protein-peptide complex structures, and systematic evaluation focused on the AlphaFold family is notably lacking. This gap necessitates a thorough evaluation of leading computational tools like AFM and CF, especially in the context of protein-peptide structure predictions using multiple quality and scoring functions. A comparative analysis of scoring functions will illuminate the relative strengths and pinpoint areas for potential enhancement in the methods applied.

In this study, to systematically evaluate the performance of prediction tools and scoring functions, we compiled a dataset of 60 native protein-peptide structures from the Protein Data Bank (PDB)[37, 38]. We used AFM to predict 1,000 protein-peptide structures, and we employed CF was employed to predict five complex structures using both TB and TF approaches. Additionally, we used AF3 was used to predict five models. This dataset served as a reference to assess the effectiveness of these three prediction tools and their scoring functions.

## Results

We conducted a comprehensive two-part evaluation and analysis of protein-peptide complex structures **(Figure 1)**. The first part focused on assessing the quality of predicted structures, while the second part aimed at analysing various scoring functions. To accomplish this, we employed two quality measures, DockQ[39] and MolProbity[40, 41], and some distinct scoring functions. The insights derived from this study have provided a nuanced understanding of the accuracy, capabilities and limitations of AFM, CF and AF3, as well as of the efficacy of each scoring function employed.

**Figure 1.**
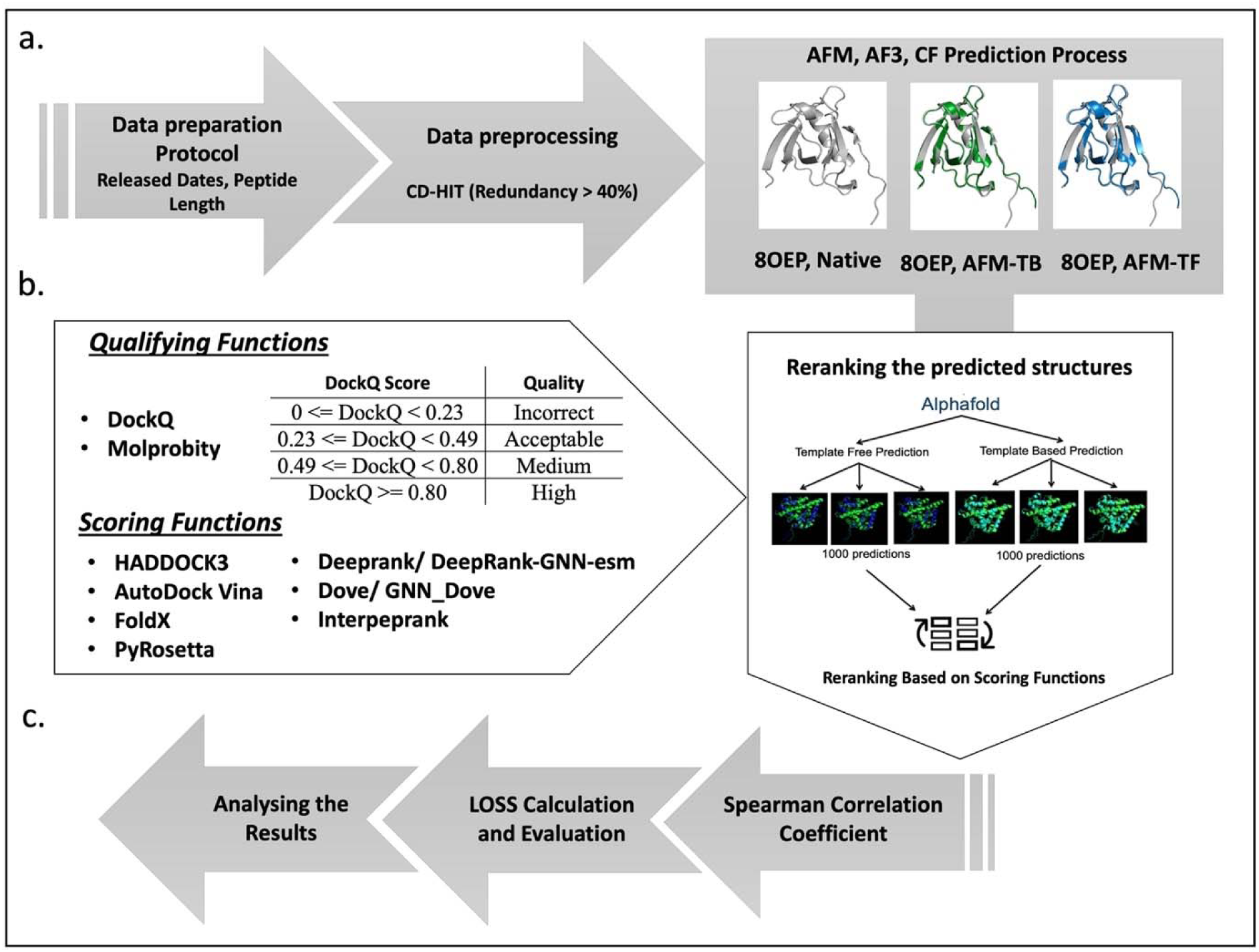
Study overview. **a.** Data preparation: The dataset comprised post-12/01/2023 protein-peptide complexes from PDB with standard amino acids in peptides of 3–50 residues, unencountered by AF’s training dataset. We removed all sequences containing nonstandard residues from the dataset. Data preprocessing: We refined the dataset using CD-HIT to eliminate sequences with over 40% redundancy, resulting in 60 diverse protein-peptide complex structures for our evaluation pipeline. All structures of sample sequences were predicted by AFM, CF and AF3. **b**. We categorized the predicted structures into template-based (TB) and template-free (TF) models based on the prediction method. We subsequently evaluated and analysed them using various quality and scoring functions. **c.** We calculated the Spearman correlation coefficient for the rankings produced by all scoring functions compared to the ranking by DockQ. We defined the loss parameter to evaluate the accuracy of each scoring function in identifying the most near-native structure from the pool.

### Quality Analysis

In this section, we provide a detailed analysis using DockQ to assess prediction accuracies against our ground-truth dataset for three leading protein structure prediction tools—AFM, CF, and AF3—across 60 samples. Additionally, we used Molprobity to diagnose structural problems in the predicted models. For each protein, AFM produced 1,000 predictions, while CF and AF3 produced 5. We compared the predicted complex structural models with their corresponding native structures using the DockQ metrics. We segmented the datasets into four categories based on DockQs prediction quality: High, Medium, Acceptable, and Incorrect. This classification evaluates the practicality of the predicted structures for subsequent biological and biochemical analyses. We used MolProbity to assess the quality of complex structures in terms of their potential issues without knowing the native structures.

#### DockQ

DockQ is a scoring tool used to evaluate the quality of protein-protein docking models by comparing predicted structures with reference (native) structures. It provides a single metric that combines interface root-mean-square deviation (iRMSD), ligand RMSD (LRMSD), and fraction of native contacts (Fnat), facilitating the assessment of docking accuracy and model quality[39, 42]. DockQ scores, ranging from 0 to 1, assess the accuracy of predicted models against native PDB structures. Scores above 0.23 are considered acceptable by the CAPRI standards[43] **(Figure 1)**. These scores indicate the quality of the interaction interface between the protein and the peptide relative to the native structure. **Figure 2** illustrates the evaluation and analysis outcomes for TB and TF methods, depicting the qualities of the top-ranked models by AFM, CF and AF3.

**Figure 2.**
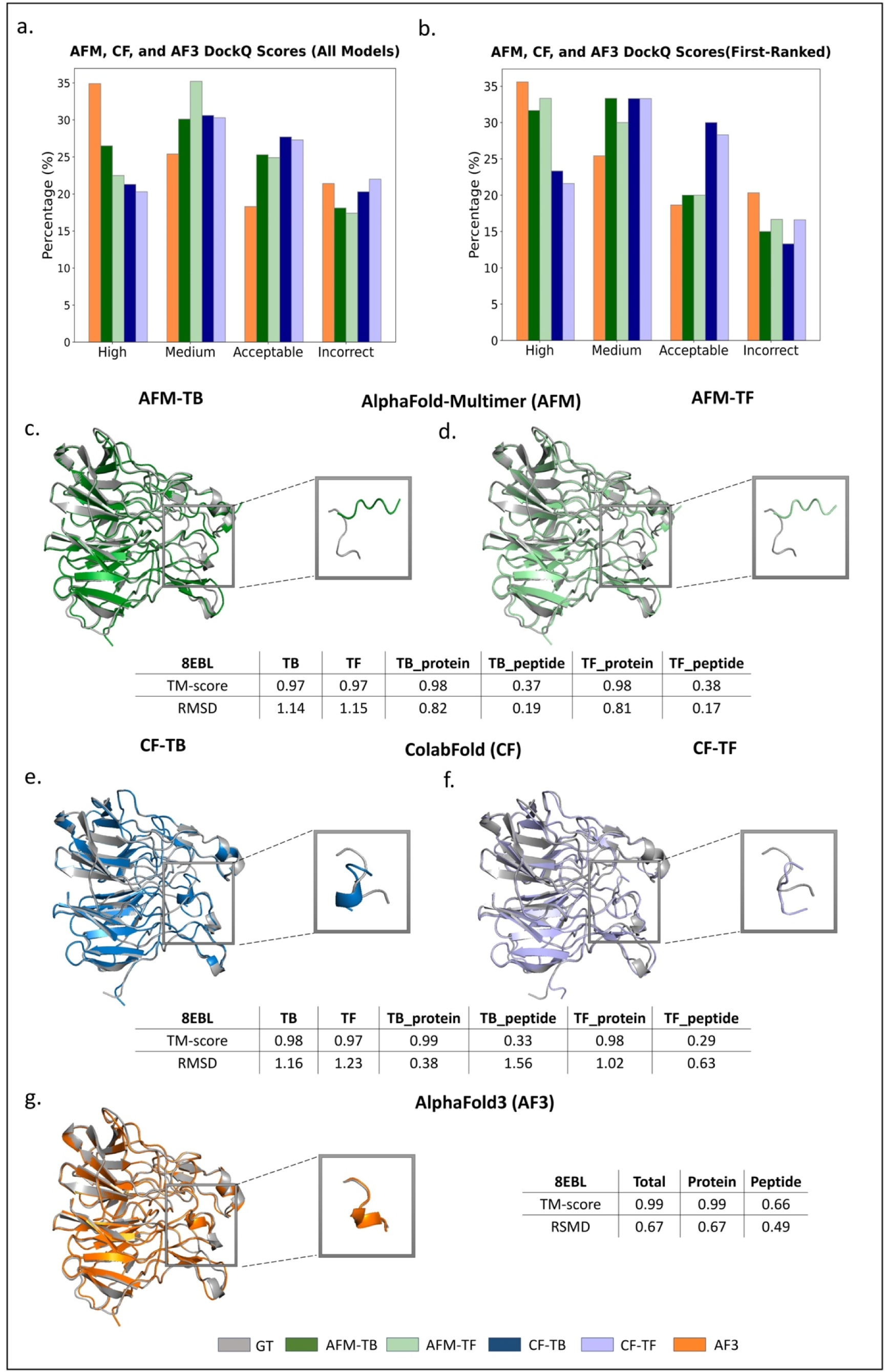
Evaluation of predicted protein-peptide complex structures using AFM, CF and AF3. **a.** Percentage of predicted structures by all three tools in different categories of DockQ. **b.** Percentage of first-ranked predicted structures by all three tools in different categories of DockQ. **c., d., e., f.** Comparative analysis of protein-peptide complex predictions for sample 8EBL by AFM and CF, using TB and TF methods, respectively, assessed against the ground truth (GT) structure with TM-score and RMSD metrics. **g.** Analysis of predicted structure with AF3 against the GT.

With AFM, the TF approach primarily produced structures in the Medium category **(Figure 2a)**. The TB approach generated more high-quality predictions than TF, demonstrating the critical role of template information in achieving optimal prediction accuracy. AF3 outperformed both AFM and CF in predicting more accurate structures, with approximately 35% of the total samples falling into the High category. CF’s performance, on the other hand, was more evenly distributed between the TB and TF approaches. The TF method exhibited a slight decline in the High category performance, achieving 20.3% vs. 21.3% in TB. Yet, it maintained similar percentages across the remaining categories (30.3% TF vs. 30.6% TB Medium, 27.3% TF vs. 27.7% TB Acceptable, and 22.0% TF vs. 20.3% TB Incorrect). This pattern suggests a minor difference between TB and TF approaches within CF.

The side-by-side comparison of AFM and CF unveils significant insights into their predictive strengths. AFM demonstrated superior performance in predicting high and medium quality structures compared to CF (26.5% AFM-TB vs. 21.3% CF-TB and 22.5% AFM-TF vs. 20.3% CF-TF in the High category, 30.1% AFM-TB vs. 30.6% CF-TB (almost similar) and 35.2% AFM-TF vs. 30.3% CF-TF in Medium category), which indicates that the convenience of using CF has a cost. These results underscore the importance of selecting AFM to predict high-quality structures over CF if possible. Nevertheless, the AFM’s performance is significantly surpassed by that of AF3, which offers both high quality and convenience.

In protein structure prediction, importance is often placed on the quality of the first-ranked structures selected by the prediction tools without knowing the native structures, as these predictions are usually used in subsequent biological analyses and functional studies. Consequently, we compared the quality of first-ranked protein structure predictions by AFM and CF by employing DockQ scores across TB and TF approaches (**Figure 2b**). For AFM, TF slightly outperforms TB in the High category (33.3% TF vs. 31.6% TB). Interestingly, although TB’s average performance is better than TF for all prediction models, the 1,000 models predicted by TF are expected to be more diverse than TB, and AFM has a strong capacity to select the best model from the pool. A substantial consistency between TB and TF approaches is demonstrated by CF, with a near-equal distribution of High and Medium quality predictions observed, although its TB approach is slightly better. This may be because CF predicted a small number of models in both TB and TF, resulting in little diversity in the model pool. AF3 generated the most high-quality structures, with 35.6% of the first-ranked models falling into this category, followed by 25.4% in the Medium category.

**Figures 2cd, 2ef** and **2g** illustrate the first-ranked protein-peptide structures predicted by AFM, CF and AF3, respectively, using both TB and TF approaches for the same sample (8EBL used as an example). To analyse the prediction accuracy of each method, we calculated global superposition metric template modelling score (TM-score)[44] and RMSD[45] metrics for the entire complex structure, and the protein and peptide components separately. Based on the results, the accuracy of protein-peptide complex structure predictions by both AFM/AF3 and CF was High, with TM-scores values around 0.97–0.99 and RMSD values around 0.67–1.23 in both TB and TF approaches. For the protein the TM-score values were within the range of 0.98 to 0.99. However, RMSD values varied, with the lowest associated with AF3 predictions and the highest recorded for AFM using the TB approach. The predicted peptide structure has worse quality than the protein, with TM-scores decreased to 0.37/0.66/0.33 for AFM-TB, AF3 and CF, respectively. AF3 outperformed both AFM and CF in terms of TM-score for the predicted peptide.

The distribution of all DockQ parameters, including Fnat, iRMSD, ligand LRMSD, DockQ, and intersection over union (IOU), was analysed in Supplementary Figs. S1-S5. While AF3 has the highest proportion of high-quality scores (DockQ ≥ 0.80), its average prediction accuracy, as indicated by the median and quartiles in Supplementary Fig. S4, is less favourable than other tools. Additionally, the radar plot in **Figure 3a** analysed the median values of the DockQ parameters. The superior performance of AFM-TB and AFM-TF in producing high-quality and consistent protein-peptide docking predictions is underscored by this analysis.

**Figure 3.**
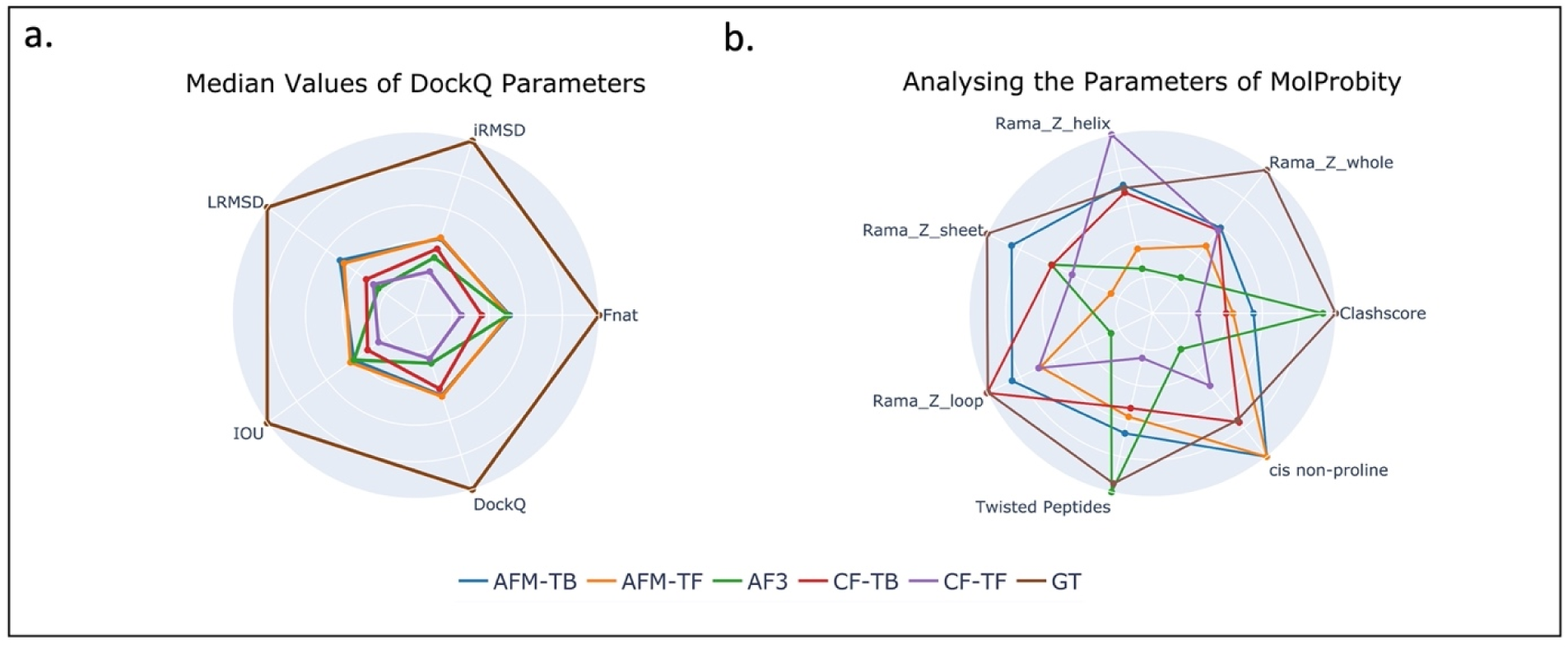
Radar plots for the prediction-accuracy strengths and the structure-quality strengths of various methods on the first-ranked predicted protein-peptide models. In each metric, the best performer takes the position of the most outer circle, the worst one takes the most inner circle, and the others take positions using linear scales of the metric. **a.** The radar plot of median values for Fnat, iRMSD, LRMSD, IOU, and DockQ parameters across five methods along with GT, which has a score of 1 for all metrics. AFM-TB and AFM-TF demonstrate high accuracy and consistency in predictions, while AF3 performs moderately. CF-TB and CF-TF have lower prediction quality, highlighting the superiority of AFM models. **b.** The radar plot of MolProbity metrics. The Clashscore is assessed by its median value of the MolProbity parameter, the smaller the better. Rama_Z scores are assessed by their root mean square values of the MolProbity parameters, also the smaller the better. GT shows the best overall performance, with the lowest RMS values across Rama_Z metrics. AF3 struggles with higher RMS values but performs well in Twisted Peptides and Clashscores. CF-TB and CF-TF excel in specific areas. GT remains the most reliable for structural quality, with AF3 and CF-TF showing targeted strengths and weaknesses.

#### MolProbity

MolProbity is a popular validation tool for assessing protein structure quality through stereochemistry and atomic interactions. It examines amino-acid residue conformations, angles, and bond lengths, pinpointing deviations from standard values. In this study, we used the MolProbity score, along with analyses of twisted peptides and cis non-proline conformations, were used to evaluate protein-peptide complex structures. In **Figure 4a**, the framework for our analysis is provided by MolProbity scores, which offer a detailed look at the quality of structural predictions from AFM, CF, and AF3. These scores inform us of the accuracy distribution within the TB or TF approach (for AFM and CF) and AF3, with higher structural quality indicated by lower MolProbity scores. The MolProbity score combines the Clashscore, rotamer, and Ramachandran evaluations into a single comprehensive score[41].

**Figure. 4.**
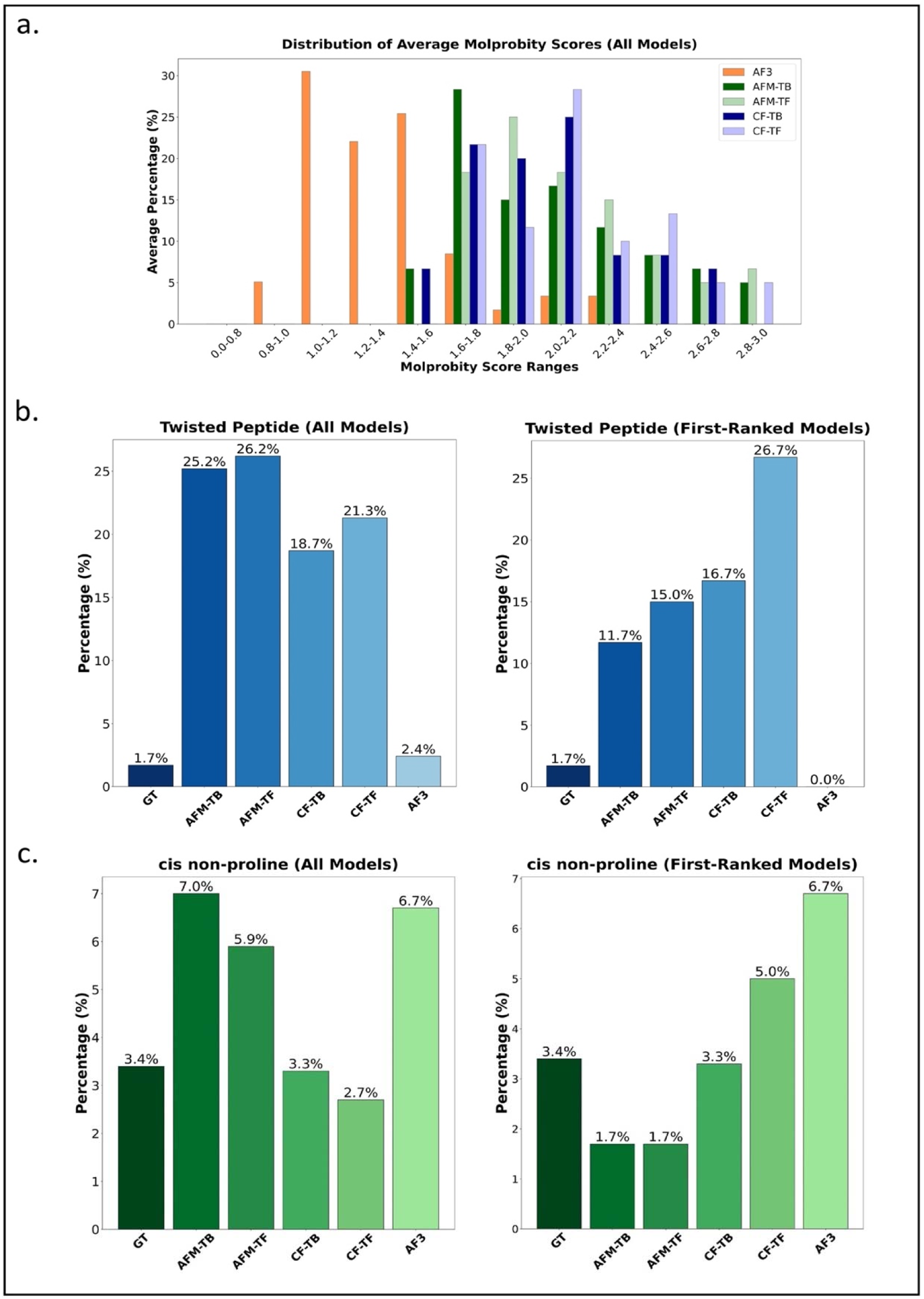
Molprobity, twisted peptides, cis non-proline peptide evaluation. **a.** MolProbity scores for AFM and CF using TB and TF approaches and AF3, higher structural quality indicated by lower MolProbity scores. **b., c.** Percentages of twisted peptides and cis non-proline conformations observed in the predicted structures for all samples and the first-ranked model, using AFM, CF and AF3. Predicted structures with fewer twisted peptides and cis non-proline conformation, or those closer to the GT models, are preferred.

The AF3 model demonstrates superior performance, with 30.5% of its predictions falling within the 1.0–1.2 range. Additionally, 22.0% and 25.4% of predictions fall within the 1.2–1.4 and 1.4–1.6 ranges, respectively. AF3’s minimal representation in high MolProbity ranges illustrates its effectiveness in generating high-quality structures. In contrast, the AFM-TB and AFM-TF models show a broader distribution of MolProbity scores. AFM-TB’s highest percentage (28.3%) falls within the 1.6–1.8 range, while AFM-TF has a significant concentration (25.0%) in the 1.8–2.0 range. Both models also have a substantial presence in the 2.0–2.2 range, indicating that they generate lower-quality structures more frequently than AF3.

The CF-TB and CF-TF models display even more varied distributions, with notable peaks in higher MolProbity score ranges. CF-TB has significant proportions in the 1.6–1.8 (21.7%) and 2.0–2.2 (25.0%) ranges, while CF-TF peaks in the 2.0–2.2 range (28.3%), with slightly worse quality than CF-TB. These models exhibit substantial percentages in ranges beyond 2.2, indicating a higher incidence of lower-quality prediction. Overall, AF3 consistently produces more high-quality predictions, while AFM-TB and AFM-TF show broader quality distributions. The Colab models display the most varied distributions, with the poorest overall prediction quality.

Significant differences between computational prediction methods were revealed by our comparative evaluation of twisted peptides within protein-peptide complexes. By considering all models of all samples **(Figure 4b)**, only 1.7% of the ground truth (GT) structures contain twisted peptides. AF3, with a low percentage of 2.4%, performs closest to the GT, suggesting high quality overall with a much smaller percentage than other methods. The percentage for CF-TB is 18.7%, which is large but still smaller than the percentages for AFM-TB, AFM-TF, and CF-TF, which range from 21% to 26% (specifically, 25.2%, 26.2%, and 21.3%, respectively). It is also important to note that AFM-TF and AFM-TB each have 1,000 predictions, whereas AF3 and CF each have only five models in their predictions, potentially influencing the results.

For the first-ranked models, comparing against the baseline, i.e., 1.7% of the GT structures with twisted peptides, AFM-TB has 11.7%, followed by AFM-TF and CF-TB with 15.0% and 16.7%, respectively. The lowest quality first-ranked models are produced by CF-TF, with the highest percentage at 26.7%. Interestingly, a 0.0% is observed for AF3, implying that no twisted peptides were identified in first-ranked models. We note that other than those generated by AFM-TF, first-ranked models have fewer twisted peptides than all other models on average.

We conducted a comparative analysis on the presence of cis non-proline peptides (**Figure 4c**, with an example shown in Supplementary Fig. S6). As a baseline for evaluation, 3.4% of the GT structures contain cis non-proline peptides. Models by AFM-TB and AFM-TF contain higher percentages of cis non-proline peptides, at 7.1% for all models (1.7% for the first models) and 5.9% for all models (1.7% for the first models), respectively. CF-TB predicted 3.3% for both all models and the first models, closely aligning with the GT structures of 3.4%, while CF-TF models have a lower percentage of 2.7% for all models (5.0% for the first models). AF3 models have more cis non-proline peptides, with a rate of 6.7% for both all models and the first models. According to our evaluation, all observed cis non-proline peptides occurred within the proteins, without any occurrence in the peptides.

All the distributions of Ramachandran scores[46] obtained from the MolProbity tool for first-ranked models are shown in Supplementary Figs. S7-S10, illustrating the distribution of Rama-Z-whole scores, Rama-Z-helix scores, Rama-Z-sheet scores, and Rama-Z-loop scores, respectively. Rama-Z scores ranging from 0 to ±1 indicate that the dihedral angles are very similar to those found in GT structures. Rama-Z scores at ±1 to ±2 show that the models slightly deviate from the normal dihedral angle distributions but are still acceptable. Rama-Z scores beyond ±2 mark significant deviations from the expected values, suggesting potential issues with the protein models, such as unusual conformations that could be errors. All the methods delivered similar performance in these Rama-Z scores, significantly worse than the GT. In particular, the GT has a significantly different distribution of Rama-Z-helix scores from other methods, with more models with Rama-Z-helix scores in the range 0 to ±1 than other methods (mostly negative values for GT while mostly positive values for other methods). This suggests that the helices generated by all models are not quite protein-like. AF3 has better Rama-Z-helix and Rama-Z-sheet scores, but worse Rama-Z-loop scores than other methods. Additionally, the distribution of Clashscore values for first-ranked models across different protein modelling methods is shown in Supplementary Fig. S11. AF3 shows the least clashes among all other models, close to the GT, while other methods have significantly higher clashes. In **Figure 3b**, the radar plot related to the MolProbity parameters displays the strengths of each model for all first-ranked models across all protein-peptide samples. AF3 demonstrates the best overall geometry for twisted peptides and Clashscores.

**Figure 5** presents example protein-peptide structures predicted by various methods together with their MolProbity metrics. **Figure 5a** displays the hydrogen bonding interactions between the protein and peptide in the 8HLO sample. It demonstrates that the hydrogen bonding in the native structure is stronger than predicted structures by AFM-TB, AFM-TF and AF3. In **Figure 5b**, the cis non-proline conformations in the peptide portion of the 8EBL sample are depicted. As illustrated, the AFM-TB models exhibit a cis non-proline conformation, indicated by ω° values (a cis peptide is characterized by an ω angle between −30° and +30°, whereas a trans peptide is defined by an ω angle greater than +150° or less than −150°[47]), whereas this conformation is absent in the native, AFM-TF and AF3 peptide structures. In **Figure 5c**, the presence of twisted peptides (ω angles that deviate more than 30° from planar[47]) in the peptide portion of the 7Z7C sample is shown for both AFM-TB and AFM-TF peptide structures, while AF3 does not show any twisted peptide in that region. In this instance, the AFM-TB model exhibits more twisted peptides than the AFM-TF models, despite the benefit of using templates for predicting the structures. In **Figure 5d**, the clashes in the peptide portion of the 8AFI sample are depicted.

**Figure 5.**
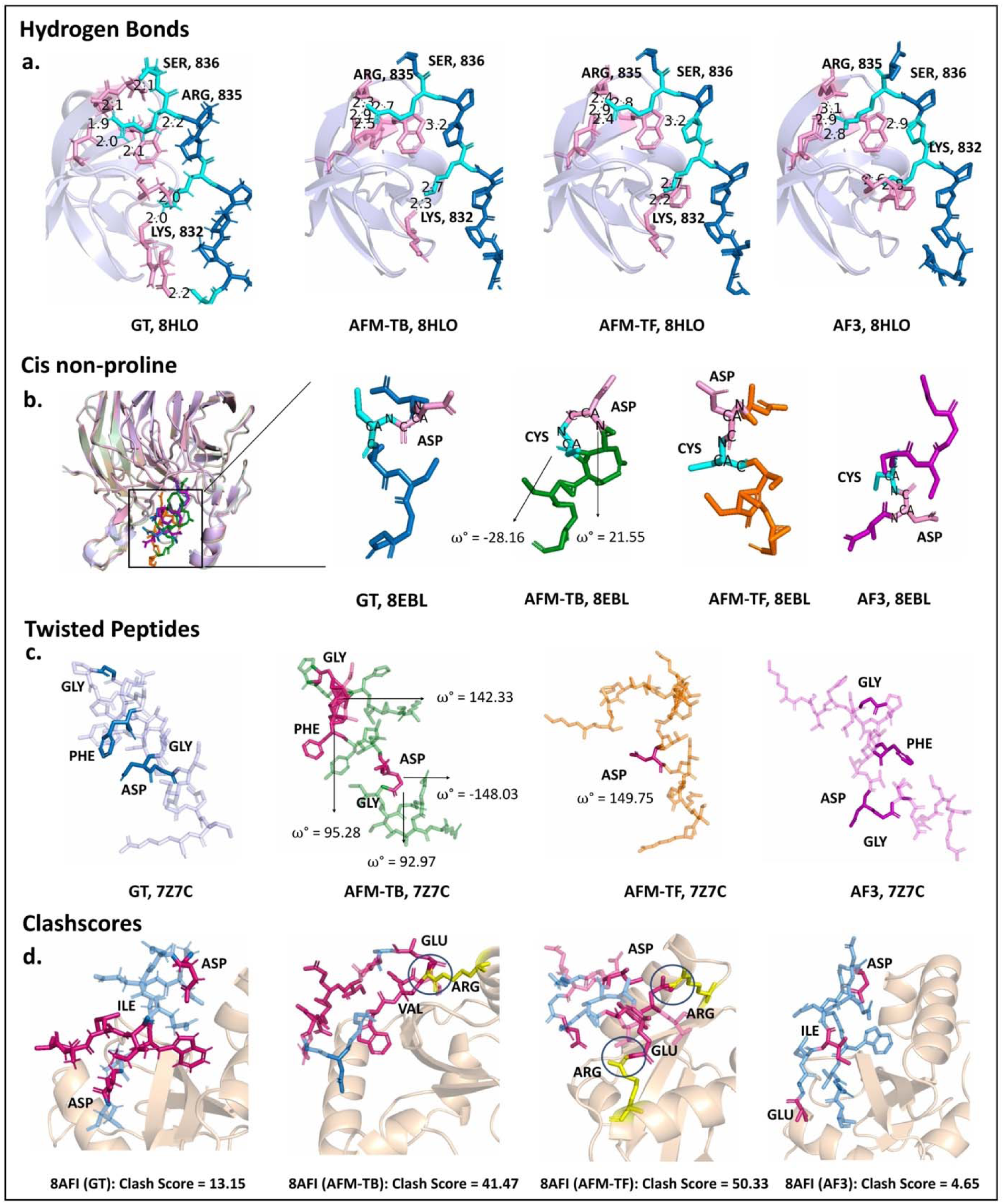
Hydrogen bonding, cis non-proline, twisted peptides, clash evaluation examples. **a.** Hydrogen bonding across protein-peptide interface of 8HLO. **b.** Cis non-proline in 8EBL. An example of a predicted structure ranked 511 was used for both TB and TF in this figure (top-ranked structures do not have any cis non-proline peptides). Corresponding residues through all structures are indicated by pink and cyan colours. **c.** Twisted peptide on peptide region of 7Z7C GT and predicted structures. Twisted peptides, marked by unusual omega (ω°) dihedral angles, deviate from standard cis (ω ≈ 0) or trans (ω≈180°) conformations. These deviations may indicate structural anomalies or prediction errors. ω angles such as 92.97°, 95.28°, −148.03°, 142.33° on TB model, or 149.75° on TF model are neither close to the cis nor the trans conformations, indicating twisted peptide bonds. **d.** The clashes on the peptide site of the complexes, along with their corresponding Clashscore for the entire complex structure. The AF3 model exhibited the fewest clashes on the peptide site in the 8AFI sample.

### Evaluation of Scoring Functions

In this section, we demonstrate the effectiveness of various scoring functions in ranking AFM-predicted protein-peptide complex structures. Using the Spearman correlation coefficient[48], we assessed the alignment of each scoring function with DockQ and other scoring functions. To further refine the analysis, all TB and TF predicted structures by AFM were consolidated to create a unique and comprehensive dataset (2,000 models for each protein-peptide complex). This approach enabled the identification of a set of common candidates, selected across different scoring functions, which potentially outperforms others in identifying near-native structures.

#### Spearman correlation matrix between scoring functions

In **Figure 6a,b**, Spearman correlation matrices are utilized to elucidate the relationships between various scoring functions under both TB and TF modelling approaches. These heatmap plots illustrate that each scoring function employs a distinct strategy and algorithm to score the protein-peptide structures in our dataset. In both TB and TF models, the AlphaFold-Multimer scoring function (AFM-Score) demonstrate moderate correlations with tools such as PyRosetta, FoldX-Stability, and HADDOCK. However, the correlations observed among HADDOCK’s variants are surprisingly modest, suggesting subtle methodological differences despite sharing a common computational framework. The minimal correlations observed between tools may mean they are complementary and have the potential to be combined for better ranking performance. The last row in both heatmap plots displays the mean correlation values of each scoring function with all other functions, excluding itself. This metric provides insight into the overall agreement and consistency of each scoring function with the others, offering a comprehensive view of their relative performance and reliability. We observe that AFM-Score exhibits the highest mean values in both the TB and TF approaches, followed by PyRosetta and HADDOCK scoring functions. This may indicate that the AFM-Score captures the strengths of other scoring functions in the most comprehensive way.

**Figure 6.**
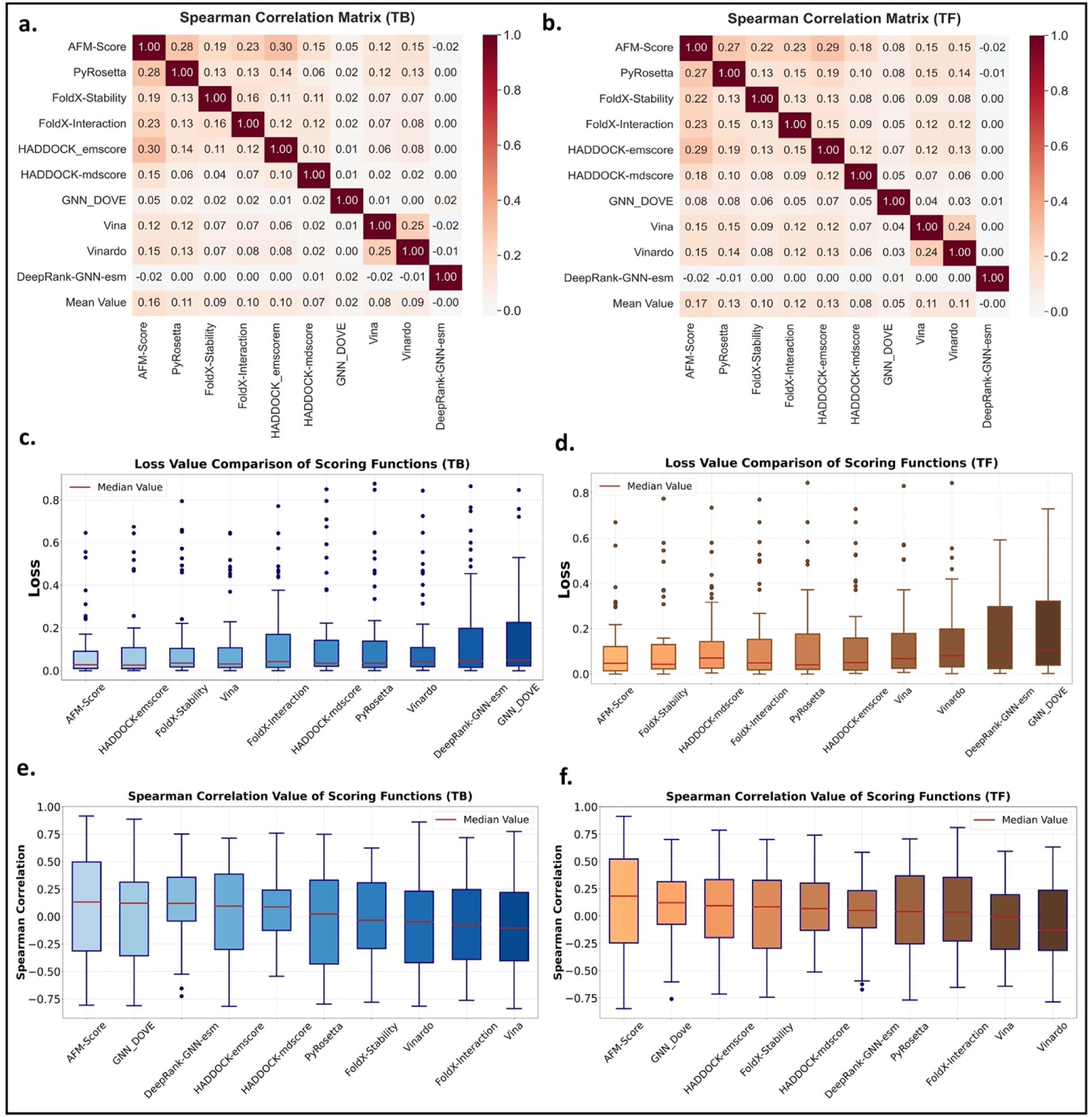
Assessing of scoring functions. **a., b.** Spearman correlation matrices on TB and TF, illustrating the similarities between each pair of scoring functions. **c., d.** Box plot distributions of LOSS values for TB and TF protein-structure predictions. These plots compare the mean and variability of AFM-Score and other scoring functions against DockQ benchmarks. **e., f.** Spearman correlation values for all scoring functions across all samples in both TB and TF models provide insight into the consistency and agreement of the scoring functions.

#### Evaluation of scoring function performance via DockQ score loss

To precisely evaluate the performance of each scoring function in finding the most near-native structure, we introduced a parameter to represent the difference between the DockQ scores of two models: (1) the top-ranked model, found by DockQ, and (2) the top-ranked model found by each scoring function. This parameter, termed LOSS, is defined as below:

*LOSS = (DockQ score of the first-ranked structure found by DockQ) - (DockQ score of the first-ranked structure found by a scoring function)*

We applied this analysis to both TB and TF methods. The results for TB and TF methods are displayed in **Figure 6c,d**, with all boxes sorted based on the mean LOSS value. The red line in each box represents the median value. The deviations in DockQ scores among various scoring functions for TB and TF predicted structures were scrutinized and quantified by the LOSS parameter. This metric provides insight into the precision of each function in ranking the top-scoring structures compared to the DockQ rankings. In **Figure 6c**, using box plots for TB protein structure predictions, the LOSS parameter across various scoring functions is examined in comparison to DockQ benchmarks on TB models. AFM-Score and HADDOCK-emscore exhibit the lowest mean LOSS values, at 0.08 and 0.09, respectively, indicating high accuracy and minimal variability. In contrast, DeepRank-GNN-esm and GNN_DOVE show the highest mean LOSS values, around 0.16, with significant variability. PyRosetta and FoldX-Stability demonstrate intermediate performance.

In **Figure 6d**, the LOSS value on TF models is illustrated. DeepRank-GNN-esm and GNN_Dove have the highest mean LOSS values of 0.17, suggesting less accurate rankings, with IQRs (Interquartile Ranges, representing the middle 50% of the data) of 0.27 for DeepRank-GNN-esm and 0.28 for GNN_DOVE, reflecting significant variability. PyRosetta and FoldX-Interaction show intermediate performance. Like in the TB case, AFM-Score still has the lowest mean LOSS value of 0.10 and an IQR of 0.10 and an IQR of 0.11. This LOSS value is higher than the one in TB (0.08), probably because the TF models are more diverse. The second-best performer is FoldX-Stability, with a mean LOSS value of 0.11 and an IQR of 0.10. HADDOCK-mdscore has the third-best performance, exhibiting the lowest mean LOSS values of 0.12. As shown in **Figure 6c,d**, AFM-Score has excelled in structure prediction and aligning with the GT samples based on the DockQ score. However, it has limitations, with eight outliers in either TB or TF models. These outliers include 7udl, 7yue, 7ue2, 8ebl, 8ahs, and 8c2p in both TB and TF, as well as 8fk3 and 7zx4 in TB, and 8b58 and 8dgm in TF. Outliers in AFM-Score predictions can be attributed to factors such as protein complexity, data quality, model limitations, dynamics, and errors in reference structures. Identifying these outliers aids in refining predictive models.

This comprehensive analysis underscores the diversity in performance among the scoring functions, revealing that while several tools perform well, their consistency differs. The variation in the LOSS values, as indicated by their IQRs and outliers, reflects the complexities of AFM protein structure prediction. Nevertheless, the comparable median LOSS values of several scoring functions to AFM-Score illustrate that multiple methods have the potential to offer similar predictive capability, though with varying levels of dependability. For example, while the medians of the three scoring functions, PyRosetta, FoldX-Stability and HADDOCK-emscore in the TF plot are lower and better than those of AFM-Score, the variability of AFM-Score remains better compared to those three. The details of the scoring functions values are shown in Supplementary Tables S2 and S3.

It is interesting to note that AF3’s ranking of the five models may not indicate their relative performance, as shown in Supplementary Fig. S12. AF predicts many nearer-native structures. The number of top 10 ranking positions for predicted structures with AF3 is calculated for each model as follows: First Rank—6, Second Rank—12, Third Rank—13, Fourth Rank—8, Fifth Rank—10. However, the third- and second-ranked models of AF3 perform better than its first-ranked positions based on these numbers. This suggests that using all five AF3 models instead of just the first-ranked one may be helpful in practical applications.

#### Spearman correlation between scoring function ranking and DockQ ranking for all models

To further evaluate the overall performance of each scoring function, we analysed all related Spearman correlation coefficient[48] values for each scoring function compared with DockQ rankings. In TB modelling **(Figure 6e)**, AFM-Score led the scoring functions with a median Spearman correlation of 0.13 and a range from −0.80 to 0.91. A strong performance was also exhibited by DeepRank-GNN-esm and GNN_DOVE, with medians around 0.12. In contrast, more consistent but moderate performance was displayed by HADDOCK scores. Lower, inconsistent results were exhibited by PyRosetta, FoldX-Stability and FoldX-Interaction. High variability was shown in the results of Vina and Vinardo.

In TF modelling **(Figure 6f)**, strong adaptability is indicated by AFM-Score, which excels with a median Spearman correlation of around 0.18 (higher than that in TB, probably due to a more diverse model pool in TF). The second performer is GNN_DOVE with a 0.12 median value. Moderate effectiveness and stability are shown by HADDOCK scores. Challenges in non-template settings are highlighted by FoldX-Interaction and Vina, which struggle with near-zero or negative medians. Conversely, Vinardo and DeepRank-GNN-esm demonstrate modest performance. Overall, robust performance is given by AFM-Score and GNN_DOVE. All related details and information about these two Spearman correlation plots are shown in Supplementary Tables S4 and S5.

Additionally, the percentage of positive Spearman correlations across all scoring functions on both TB and TF models are illustrated in Supplementary Fig. S13. Although some tools show positive correlation and better median Spearman correlation coefficients, they all fail to align well with the ground-truth DockQ ranking. For instance, despite a good overall Spearman correlation between GNN_DOVE and DockQ, GNN_DOVEs performance with TF models in identifying the best model for each protein-peptide complex is weaker than most other tools.

In TB and TF models, the highest median Spearman correlations are achieved by AFM-Score, showcasing AFM-Score’s adaptability across diverse conditions. GNN_DOVE and DeepRank-GNN-esm display strong performances in TB models, but DeepRank-GNN-esm effectiveness is slightly reduced in analysing the predicted structures without templates, indicating a reliance on structured data. HADDOCK demonstrates a consistent scoring approach for both TB and TF models. In less constrained environments, underperformance is highlighted by FoldX-Stability and FoldX Interaction, showing their limitations. The variability of Vina and Vinardo is consistent in both contexts; however, their unpredictability is more pronounced in TF environments, emphasizing the associated risks. Overall, while some scoring functions adapt well across different frameworks, others depend significantly on the presence of templates in the prediction process, impacting their scope of application.

Because the strong performance of the scoring functions—AFM-Score, FoldX-Stability and HADDOCK-mdscore—was demonstrated for both TB and TF models, we selected these for further evaluation. In **Figure 6c,d**, AFM-Score displayed superior overall performance in the TB models, albeit with significant variability. Conversely, FoldX-Stability and HADDOCK-mdscore exhibited higher LOSS values and wider spreads, indicating less robustness. Similarly, AFM-Score showed higher variability in TF models than TB models but maintained better performance than FoldX-Stability and HADDOCK-mdscore, which demonstrated higher LOSS values and more substantial spreads, underscoring AFM-Scores relative superiority despite its high variability.

We combined all TB and TF models from each protein-peptide sample and extracted the top 10 models (including both TB and TF top-ranked models) from each sample. The Venn diagram in **Figure 7a** demonstrated the overlap of top-ranked models among the scoring functions, identifying common samples ranked in the top 10 by all three scoring functions. These common samples from the three mentioned scoring functions (Com-score-3), characterized by consistently lower LOSS values, emerged as reliable candidates for near-native structures against the individual scoring functions. The intersection between FoldX and HADDOCK showed the lowest mean LOSS value of 0.01 compared to all other scoring functions, as illustrated in **Figure 7b,c**, suggesting that these two scoring functions are highly complementary to achieving a good ranking. The combinations of Com-score-3 recorded lower LOSS values, with a mean LOSS of 0.07, compared to 0.10, 0.11 and 0.12 for AFM-Score, FoldX-Stability, and HADDOCK-mdscore in TF models, and 0.08, 0.11 and 0.13 in TB models, respectively. As depicted in **Figure 7d,e**, adding the PyRosetta scoring function to the combination, Com-score-4, with a mean value of 0.06, shows improved performance compared to the combination of three scoring functions. Both combinations performed better than the other scoring functions.

**Figure 7.**
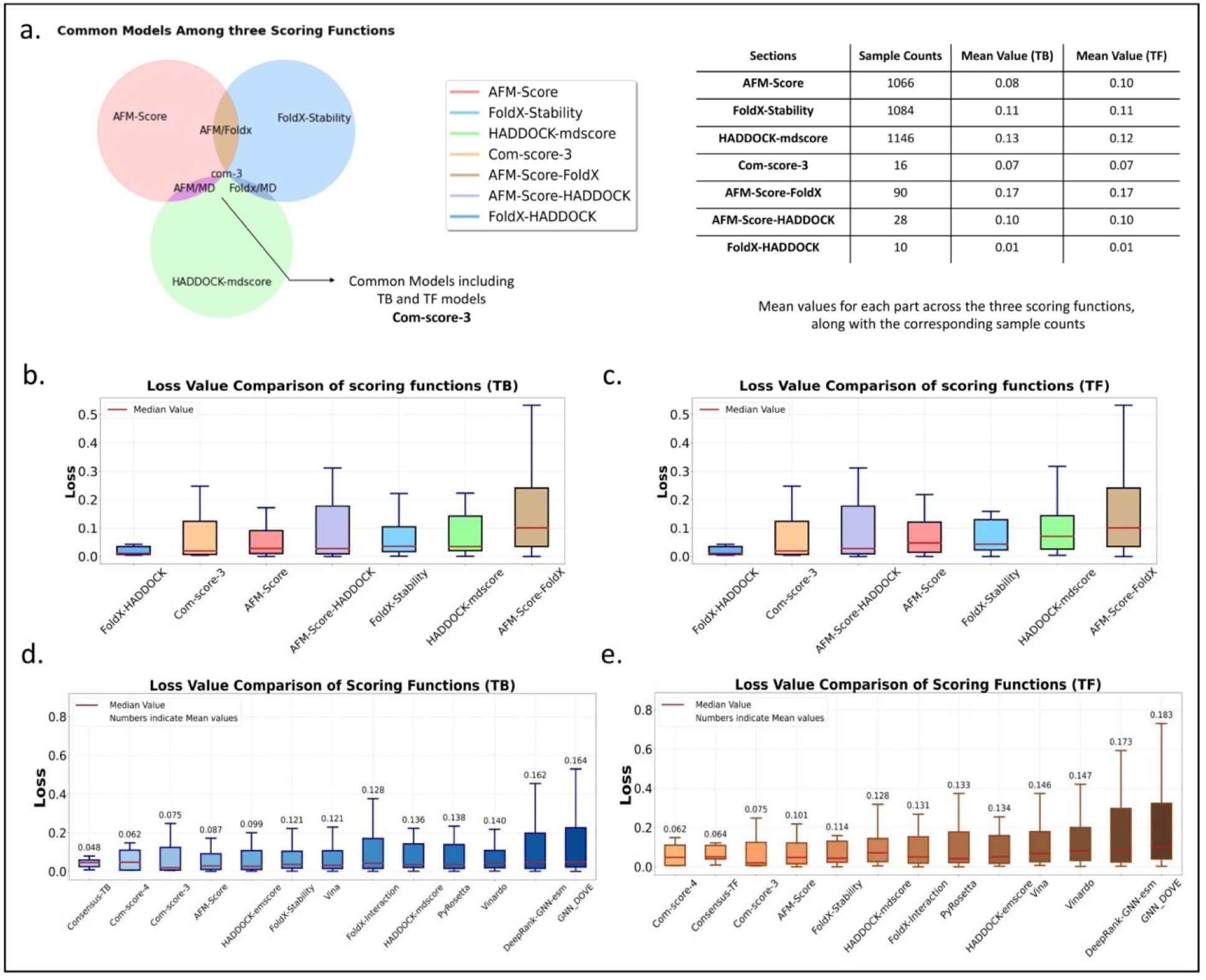
Evaluation and comparison of scoring functions for predicting near-native structures. **a.** Venn diagram illustrating the intersection of three scoring functions: AFM-Score, FoldX-Stability, and HADDOCK-mdscore. The ‘com-3’ region indicates the common samples region representing the intersection of the three scoring functions. **b., c.** Box plots for TB and TF models were analysed using the three scoring functions and common samples. In both cases, common samples (Com-score-3) demonstrate lower mean LOSS values, indicating higher accuracy in predictions**. d., e.** The intersection region among the four scoring functions, including all previous functions along with PyRosetta, was also evaluated against all other scoring functions in TB and TF models. This evaluation is represented as Com-score-4 in the plots. Both consensus-TB and Consensus-TF approaches performed better than any individual scoring function. The consensus method produced more consistent results and less variability in TB and TF models.

We also explore the potential of the consensus approach, which integrates multiple models to achieve a more reliable and accurate ranking[49]. This approach leverages the strengths and compensates for the weaknesses of individual models, leading to a more robust prediction of protein structures. For this purpose, we selected eight protein-peptide complex samples from the TB and TF datasets separately: 7wqq, 7xty, 8ese, 8ia5, 8i3g, 7prx, 8cir, 7r2m. These samples were obtained from different quartiles of the LOSS values distribution to ensure a diverse performance representation. Each of these protein-peptide complexes has 1,000 predicted structures in both TB and TF approaches. We calculated the DockQ score for each of these 1,000 models against all other 999 structures in the pool. Finally, we added up all these DockQ scores and extracted the sample with the maximum value of this sum. This consensus approach, indicated by Consensus-TB and Consensus-TF on their corresponding plots, capitalizing on the agreement among different scoring functions, exhibited superior performance, as evidenced by the box plots (**Figure 7d,e**). Notably, this approach showed the smallest spread on TB models and the lowest mean LOSS values on both TB and TF models, indicating high accuracy and reliability. The analysis highlights the advantage of using a consensus approach to identify near-native structures, showing that samples highly ranked by multiple scoring functions exhibited superior performance.

## Discussion

Our research thoroughly examined AFM, CF and AF3, targeting their accuracy, protein-like properties, and dependability in predicting protein-peptide complex structures—a research area notably lacking in systematic studies. Using benchmarks and reference structures from PDB, our work provided a detailed and comprehensive evaluation of the AlphaFold tool family and their applicable scoring functions through widely adopted quality functions, including DockQ and MolProbity, across both TB and TF methodologies.

Our analysis shows that AFM excels in TB predictions with templates but has moderate TF precision in the prediction pool, but TF outperforms TB in the first-ranked models. CF, while less accurate than AFM, is versatile and consistent in TB and TF, improving slightly with templates. AF3 generated high-quality structures for more protein samples; however, the AF3 medium DockQ score of all protein samples is significantly less than that of AFM. This highlights the importance of generating a deep pool for achieving high accuracy. As AF3 only provides five models for each protein, the ongoing community initiatives to reproduce AF3 as an open-source tool are meaningful. Such initiatives may not reach the performance of in-house AF3, but it is likely to deliver a tool with higher accuracy than AlphaFold2, and with a deep pool the tool can achieve better performance than the AF3 web service.

Our analysis of MolProbity scores for AFM, CF and AF3 reveals different trends in the quality of structural prediction. The AFM-TB approach generally produces structures of moderate quality, and the model quality remains commendable in the TF settings, even in the absence of template data. CF, on the other hand, consistently generates structures with medium quality across both TB and TF methods, with a marginal enhancement in the quality of structures when template data is used. The AF3 model demonstrates superior performance compared to the other two tools. In exploring specific cases like twisted peptides and cis non-proline residues, we note that AF3 predicted structures with significantly fewer twisted peptides than AFM or CF. However, in predicting structures with cis non-proline peptides, both AFM and CF were slightly better than AF3.

It should be noted that all the evaluated structures on AF3, including those with cis non-proline peptides, have these conformations located on the protein part rather than on the peptide site. The analysis of hydrogen bonds in the GT and predicted structures revealed that the native structure has stronger hydrogen bonding than the predicted models. In addition, the helices generated by any model have a different distribution from the GT. Despite using templates, some TB models showed more twisted peptides than the native and TF models. AF3 exhibited no clashes in models, while other methods produced some clashes. Considering all factors, AF3 delivered better protein-like properties than other methods, but still has room for improvement.

For ranking prediction models, AFM-Score is clearly better than other scoring functions. This may be partly because AFM-Score is developed explicitly for AlphaFold models, while other scoring functions are intended for general models. Some scoring functions, such as FoldX-Stability and HADDOCK-mdscore, also delivered good performance. The scoring functions are highly complementary to each other, as shown by weak correlations between any of them. This suggests the advantage of using a consensus approach to identify near-native structures, as common samples in both three-function (com-score-3) and four-function (com-score-4) combinations ranked highly by multiple scoring functions showed better performance with lower LOSS values. Our consensus method, which extracts the structure with the highest summed DockQ score among the structures in the pool, also demonstrated better performance compared to AFM-Score and all other individual scoring functions in both TB and TF models. Given the massive usage of AFM, it is worthwhile to design scoring functions specifically targeting predicted structures by AFM. Our study suggests the potential to refine individual scoring functions and integrate ensemble methods for improving the model selection of complex structures predicted by AFM.

## Conclusion

Our study provides a comprehensive evaluation of AFM, CF, and AF3 for predicting protein-peptide complex structures, highlighting both the strengths and limitations of each tool. AFM excels in Template-Based (TB) predictions, while Template-Free (TF) models often surpass TB models when ranked. CF shows consistent performance across TB and TF methodologies, and AF3 demonstrates the potential for generating high-quality structures, although a deeper model pool is needed to achieve greater accuracy. Analyzing structural quality metrics such as MolProbity scores, hydrogen bonding, and specific features like twisted peptides and cis non-proline residues reinforces the performance differences among these tools. Notably, AF3 delivered better protein-like properties with fewer clashes and twisted peptides, though some areas still show room for improvement. Our investigation into various scoring functions revealed the potential for optimizing structure selection through consensus approaches, combining metrics such as FoldX-Stability and HADDOCK-mdscore to improve accuracy. Future efforts should focus on refining scoring functions tailored to AFM models and enhancing ensemble methods to improve prediction reliability in protein-peptide docking.

## Methods and Materials

### Benchmark Dataset

In this study we developed a comprehensive pipeline to evaluate the quality of predicted protein-peptide structures by AFM, CF and AF3. Additionally, this pipeline assesses the efficacy of various scoring functions in accurately ranking these predicted structures, thereby assessing their capability to represent protein-peptide interactions accurately. For this purpose, we employed two quality functions alongside distinct scoring functions.

To facilitate a thorough evaluation of AFM and CF’s performance in predicting protein-peptide complex structures, we initiated the systematic preparation of a benchmark dataset. We carefully curated this dataset to include protein-peptide complex structures that feature at least one protein and one peptide chain, with a released date after 12/01/2023 (the cutoff date for the training set used by AFM). This criterion ensured the incorporation of novel structures not previously encountered by the predictive models. Furthermore, we restricted our selection to peptide sequences spanning three to 50 residues in length, excluding any sequences containing nonstandard amino acids to maintain dataset quality. Following the collection of all pertinent protein-peptide sequences, we employed the CD-HIT[50] tool to identify and remove sequences exhibiting more than 40% redundancy, thus enhancing the dataset’s diversity. This thorough selection process resulted in a dataset comprising 60 unique protein-peptide complex structures ready to incorporate into our evaluation pipeline (Supplementary Table S1).

### Model Generation

AFM models were generated using AlphaFold v2.3.1, employing both TB and TF approaches. AFM was run on NVIDIA A100 80GB PCIe GPUs. Default parameters were used for multiple sequence alignment generation to maintain consistency across predictions. For each protein-peptide sample, 1,000 complex structures were predicted using the TB method and 1,000 structures using the TF method, ensuring a comprehensive analysis. We also processed all samples using CF’s default settings to predict structures through TB and TF approaches. By default, each approach yielded five ranked models. The complex structures of all samples were predicted using the AF3 web server, which by default returned five predicted structures for each sample, regardless of the presence of templates. One of the samples (8C2P) out of the 60 in our dataset could not be predicted by AF3 due to a biological sequence restriction, resulting in a “Sequence filtering encountered” error.

For the AFM-TB models, the ‘max-template-date’ parameter was set to approximately seven days before the release date of the protein-peptide complex on the PDB website. This strategy was designed to incorporate all relevant PDB structures available in the AlphaFold dataset while explicitly excluding the target protein-peptide complex under investigation. Conversely, for TF models, the ‘max-template-date’ parameter was adjusted to ‘1,000-01-01’, effectively excluding all templates from the modelling process and ensuring a purely template-free approach. The approach employed both TB and TF modelling to thoroughly evaluate CF’s and AFM’s performance in predicting protein-peptide complex structures, deliberately omitting Amber’s relaxation step to assess only the unrelaxed, ranked models. AFM was run as below.

`Alphafold --use_gpu --data_dir=<ALPHAFOLD_DATADIR> --fasta_paths=<fasta file`

`path> --output_dir=<OUTPUT directory> --max_template_date=<max`

`template date> --model_preset=multimer –`

`num_multimer_predictions_per_model=200 --run_relax=false`

AFM models were generated using full-length sequences from the PDB SEQRES section. During preparation, we conducted a cleaning step on predicted PDB structures to remove any residues not resolved in the experimental data, aligning only residues present in both the modelled structures and the reference PDB ATOM section. This critical step, ensuring model integrity and accurate alignment with PDB sequences, was essential for precise DockQ score calculations and reliable comparison against existing PDB entries.

All structural images presented in this paper were generated using PyMOL software (Schrödinger, LLC).

### Evaluation with Quality Functions

We used DockQ and Molprobity to compare predicted complexes to native structures and to assess their quality, respectively. DockQ scores, ranging from 0 to 1, assess the accuracy of predicted models against native PDB structures. Scores above 0.23 are considered acceptable by CAPRI standards. These scores indicate the quality of the interaction interface between the protein and the peptide relative to the native structure. **Figure 1** illustrates the evaluation and analysis outcomes for TB and TF methods, depicting the qualification of the top-ranked models by AFM, CF, and AF3.

`./DockQ/scripts/fix_numbering.pl <cleaned predicted structures> <cleaned`

`native structure>`

The output of fix_numbering.pl (above command) was then used to calculate the DockQ score as follows:

`python DockQ.py <aligned predicted structures> <native structure>`

The DockQ score between all predicted structures and the native structure was calculated and considered in all analysis parts.

MolProbity is a tool for validating and analysing 3D protein and nucleic acid structures. It assesses structural quality by evaluating parameters such as Clashscores, rotamer outliers, and Ramachandran plot statistics, thereby ensuring model accuracy and reliability. To prepare the predicted structures for analysis with MolProbity, the phenix.reduce command was executed to add hydrogens, potentially flip side chains, and build missing atoms, thus correcting common structural issues and ensuring completeness and accuracy. The following commands were used to run MolProbity on the predicted structures.

`phenix.reduce -FLIP -BUILD <pdbfile> &> <pdbfile>FH.pdb`

`phenix.molprobity <pdbfile>FH.pdb`

In these commands, <pdbfile> is the input PDB file, and <pdbfile>FH.pdb is the output file where the results of the reduced operation are saved.

### Evaluation with Scoring Functions

AFM uses a scoring function named ‘predicted TM-score’ (pTM) to rank predicted structural models. The pTM score predicts the model’s similarity to the GT structure, aiming for a high TM-score match. DockQ, which compares predicted structures to native models, served as a reference for evaluating the ranking accuracy of the AFM, CF and AF3 scoring functions. All 1,000 predicted models from both TB and TF methods were reranked according to DockQ scores. To assess the ranking effectiveness of AFM-Score and CF-Score, we calculated the Spearman correlation coefficient between AFM’s rankings and those of DockQ. We employed a range of scoring functions to rank the AFM-predicted structures in both TB and TF models. We selected these scoring functions from various methods to analyse the predicted structures from different perspectives. They included PyRosetta, Foldx (Stability, Interaction), HADDOCK (emscore, mdscore), AutoDock Vina (Vina, Vinardo), GNN_DOVE, and DeepRank-GNN-esm (**Table 1**).

**Table 1.**
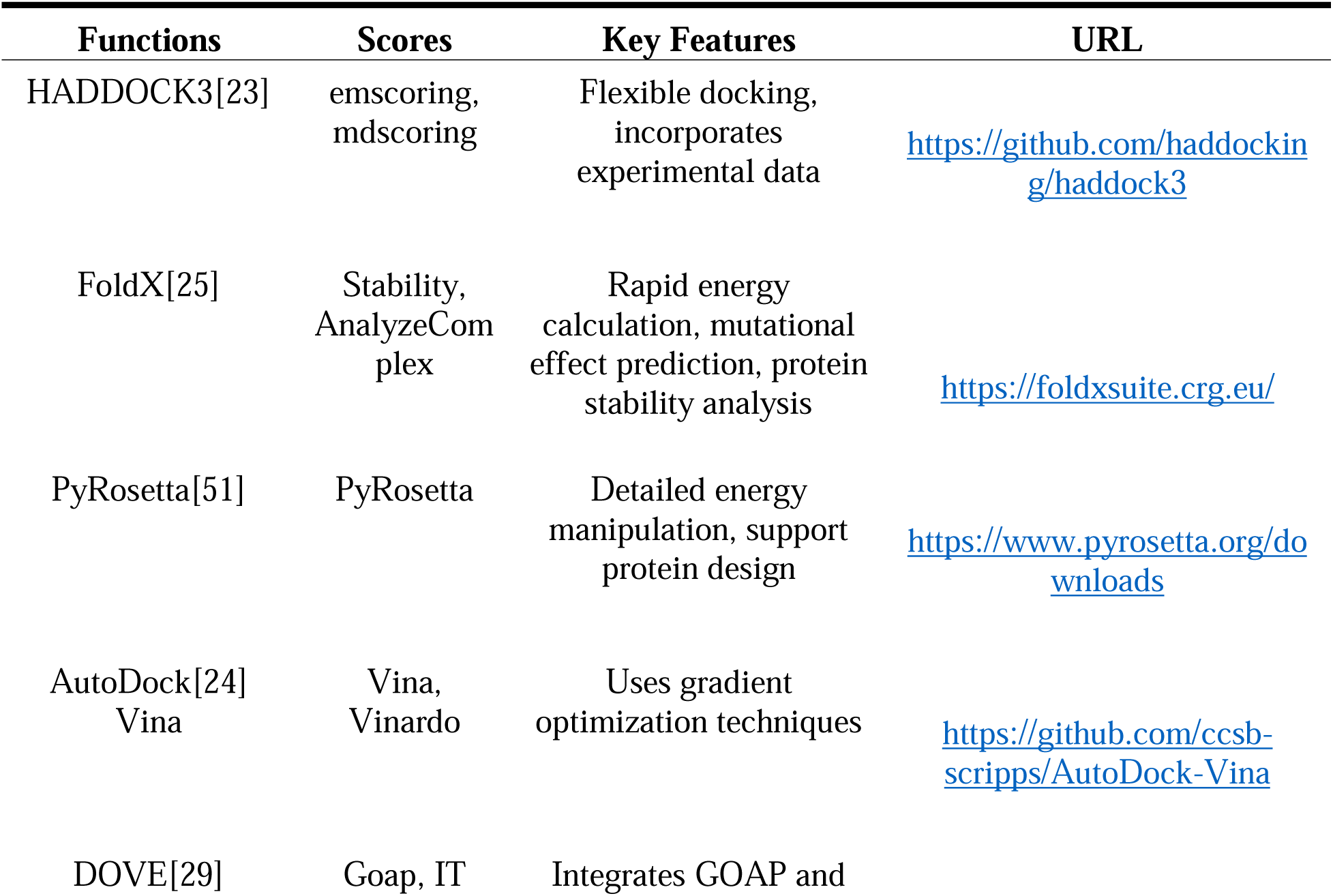
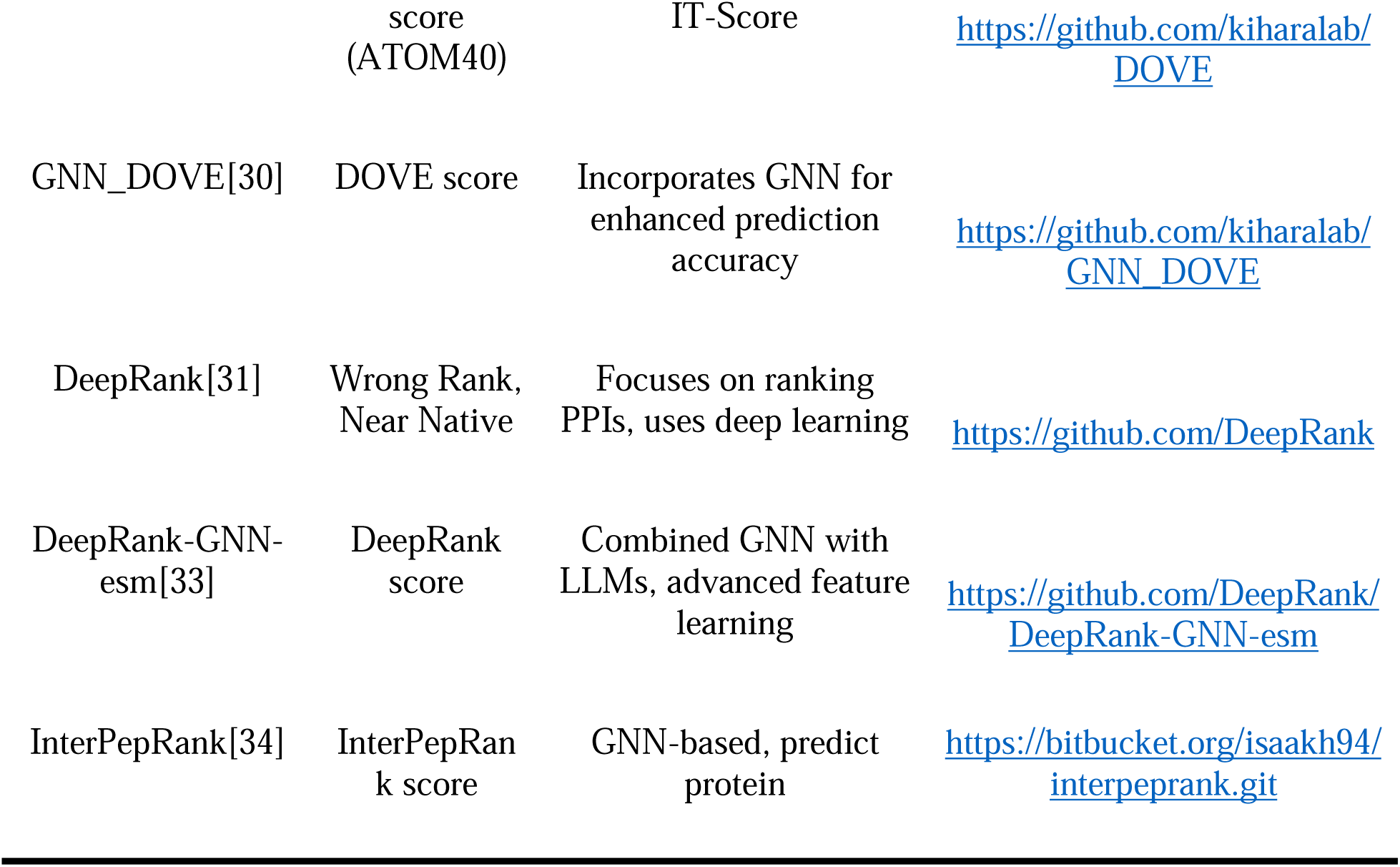

PyRosetta is a customizable toolkit for protein structure prediction and design, and we used its scoring function in this study. HADDOCK employs empirical (emscore) and molecular dynamics (mdscore) scoring for flexible docking of biomolecules. AutoDock Vina and its variant Vinardo provide an efficient scoring function based on energy calculations, considering a 5Å dimension box around the peptide part to ensure accurate modelling of critical interactions within the binding site while maintaining computational efficiency. GNN_DOVE and DeepRank-GNN-esm are two other deep-learning-based scoring functions used for accurate protein complex scoring. The Python codes for running and analysing the predicted structures with these scoring functions are available in the GitHub repository. FoldX offers both stability and interaction scoring functions to assess protein-peptide structures, and the commands used to run it are provided below. To run FoldX on a protein structure, the following command was executed sequentially.

`foldx --command=RepairPDB --pdb=example.pdb`

This command below repairs the input PDB file by adding missing atoms, optimizing the structure, and fixing any structural issues. This step ensures that the structure is complete and accurate, preparing it for further analysis.

`foldx --command=Stability --pdb= <pdbname>_Repair.pdb`

After repairing the PDB file, the above command calculates the stability of the protein structure. It estimates the free energy of folding, referred to as FoldX-Stability in this study, which can be used to assess the stability and reliability of the protein model.

`Foldx --command=AnalyseComplex --pdb= <pdbname>_Repair.pdb –`

`analyseComplexChains=A,B`

This command analyses the interactions between specified chains (in this case, chains A and B) in the repaired PDB file. It calculates various energy terms associated with the protein-peptide complex, such as binding energy, providing insights into the interaction dynamics and stability of the complex. This parameter is referred to as FoldX-Interaction in this study.

In addition to the previously mentioned scoring functions, we evaluated DOVE and DeepRank[52] were also evaluated on all 60 samples. However, due to installation issues with DOVE, which led to unsatisfactory outcomes (predominantly zero values for ATOM40), we excluded its results from our evaluation as well. DeepRank, which relies on PSSM in its algorithm, encountered difficulties in identifying necessary homologs for very short peptide sequences. As a result, it was only applicable to about 28 out of 60 samples of our dataset, leading to its exclusion from our evaluation system. Other scoring functions were also attempted for installation and testing on our dataset but could not be used. GDockScore[53] was not applicable since it only accepts peptides with more than 30 residues, while some of the peptides in our dataset had fewer than 30 residues. InterEvScore[54] encountered an error when running, and alignment only works with chains of more than 30 residues as well. PyDocks[54] download link could not be accessed.

### Evaluations

Our analysis evaluated various scoring functions using a dataset of protein-peptide complexes, comparing predicted AFM structures (TB and TF) against native PDBs for true structural fidelity. DockQ served as a reference by comparing and ranking predicted structures aligned with their native counterparts, establishing a benchmark for assessing other scoring functions. We used the Spearman correlation coefficient as the metric for this evaluation. In the below formula, if *r_i_* is the rank of the *i*^*th*^ observation in the first set, and *s_i_* is the rank of the *i^th^* observation in the second set, then *d_i_* = *r_i_*-*s_i_*.

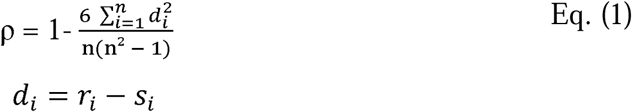

Where *d_i_* is the difference between the ranks of corresponding values, *n* is the number of observations, and 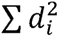 is the sum of the squares of the rank differences[55]. We used the Spearman correlation coefficient to determine the concordance level between the rankings produced by an array of scoring functions and the established DockQ benchmark. A coefficient nearing the value of 1 indicates a strong positive agreement, signifying that a scoring function’s rankings are highly consistent with those from DockQ. Such a correlation is indicative of the scoring function’s predictive reliability for the structural configurations of protein-peptide complexes.

We note that not all scoring functions provided scores for all models. AutoDock Vina’s scoring function failed for four samples—8CK5, 8ESE, 8HDJ, and 8HEP—due to time and memory limitations. Consequently, AutoDock Vina evaluated a total of 56 samples. GNN_DOVE and FoldX also failed for samples 8HEP and 8ARE in both the TB and TF approaches due to time constraints encountered during model execution. Consequently, these two scoring functions were evaluated on only 59 samples. InterPepRank’s inference on a CPU typically took two to three days per sample, indicating time inefficiency. Challenges included NaN values and uniform results due to incorrect weight loading, leading to prediction failures. Satisfactory results were not obtained for half of the samples in both TB and TF models. Due to this data inequality, we excluded InterPrepRank from our evaluations.

### Consensus Approach

This consensus approach aims to find the predicted structure with the most similarities to other structures in the pool and rank them as the best predictions. For a given protein, we calculated the DockQ score for each of 1,000 predicted structures in either TB or TF approaches against all other 999 structures in the pool. We added up all these DockQ scores and extracted the sample with the maximum value of this sum. Equations (2) and (3) show the mathematical formula of the process.

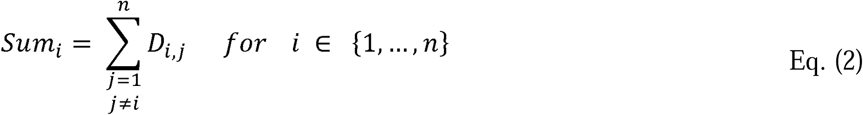

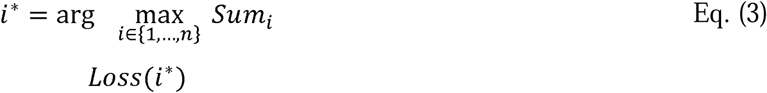

In equation (2), *D_i,j_* represents the DockQ score between the *i*^th^ and *j*^th^ predicted structure in the pool, and *i*\* shows the maximum summation value.

## Authors’ Contributions

D.X. conceptualized the study and methodology. N.M. performed the investigations, formal analyses and data curation. N.M. and J.R. implemented the validation and checking process. All authors contributed to running tools and conducting the experiments. N.M. wrote the first draft of the manuscript and created all visualizations, with J.R. providing notes on the manuscript. D.X. reviewed and edited the manuscript. Additionally, D.X. administered the project and supervised the study.

## Competing Interests

The authors declare no competing interests.

## Acknowledgements

This work is supported by the National Institutes of Health grant R35-GM126985.

## Availability of data and materials

## Dataset

The data used for this evaluation were generated during this study. Source data associated with this paper are provided in the GitHub repository (https://github.com/NeginManshour/PpEv).

## Code

The source code of this evaluation is freely available at https://github.com/NeginManshour/PpEv.

## Notes

### Competing Interest Statement

The authors have declared no competing interest.

