## Supplementary File for "Comprehensive Evaluation of AlphaFold-Multimer, AlphaFold3 and ColabFold, and Scoring Functions in Predicting Protein-Peptide Complex Structures"

**Supplementary Information**

**Supplementary Table S1.** Summary of the protein-peptide dataset. This dataset includes protein-peptide complex structures with at least one protein and one peptide chain, released after 12/01/2023. Peptide sequences range from 3 to 50 residues. Non-standard amino acids were excluded. The CD-HIT tool was utilized to remove sequences with more than 40% sequence identity.

|  | **PDB_ID** | **No. Residues (Peptide)** | **No. Residues (Protein)** | **Released Date** |
| --- | --- | --- | --- | --- |
| 1 | 7PRX | 21 | 250 | 2/8/2023 |
| 2 | 7QOX | 29 | 367 | 18/1/2023 |
| 3 | 7QWV | 18 | 148 | 2/1/2023 |
| 4 | 7R2M | 24 | 422 | 5/17/2023 |
| 5 | 7SQA | 22 | 290 | 5/10/2023 |
| 6 | 7SXF | 17 | 345 | 6/14/2023 |
| 7 | 7SXJ | 17 | 351 | 6/14/2023 |
| 8 | 7TO9 | 18 | 128 | 1/25/2023 |
| 9 | 7UDK | 12 | 172 | 3/22/2023 |
| 10 | 7UDL | 21 | 282 | 3/22/2023 |
| 11 | 7UE2 | 19 | 304 | 3/29/2023 |
| 12 | 7UI8 | 23 | 128 | 4/5/2023 |
| 13 | 7UW2 | 13 | 240 | 3/15/2023 |
| 14 | 7V1A | 19 | 301 | 6/14/2023 |
| 15 | 7WQQ | 13 | 265 | 1/25/2023 |
| 16 | 7X8G | 4 | 160 | 1/18/2023 |
| 17 | 7XFG | 16 | 102 | 4/12/2023 |
| 18 | 7XTY | 11 | 95 | 5/17/2023 |
| 19 | 7XV1 | 29 | 120 | 6/14/2023 |
| 20 | 7Y8F | 8 | 260 | 4/26/2023 |
| 21 | 7Y9C | 13 | 275 | 4/12/2023 |
| 22 | 7YUE | 15 | 253 | 2/8/2023 |
| 23 | 7Z5L | 21 | 30 | 3/22/2023 |
| 24 | 7Z6U | 14 | 313 | 2/1/2023 |
| 25 | 7Z7C | 22 | 162 | 3/29/2023 |
| 26 | 7ZAX | 16 | 133 | 6/7/2023 |
| 27 | 7ZW4 | 25 | 303 | 5/31/2023 |
| 28 | 7ZX4 | 21 | 364 | 3/29/2023 |
| 29 | 8A68 | 9 | 236 | 4/26/2023 |
| 30 | 8AFI | 24 | 115 | 5/10/2023 |
| 31 | 8AHS | 26 | 149 | 3/22/2023 |
| 32 | 8ARE | 5 | 525 | 2/22/2023 |
| 33 | 8B58 | 11 | 211 | 2/1/2023 |
| 34 | 8BFT | 20 | 343 | 2/22/2023 |
| 35 | 8BR9 | 38 | 231 | 3/8/2023 |
| 36 | 8C2P | 24 | 480 | 4/26/2023 |
| 37 | 8C3H | 8 | 124 | 2/8/2023 |
| 38 | 8CCW | 9 | 282 | 5/10/2023 |
| 39 | 8CIR | 16 | 347 | 3/1/2023 |
| 40 | 8CZ9 | 8 | 110 | 5/31/2023 |
| 41 | 8CZK | 6 | 419 | 1/25/2023 |
| 42 | 8D51 | 22 | 103 | 6/14/2023 |
| 43 | 8D7P | 18 | 301 | 5/17/2023 |
| 44 | 8DGM | 20 | 258 | 6/7/2023 |
| 45 | 8DVL | 32 | 633 | 3/29/2023 |
| 46 | 8DWK | 46 | 299 | 2/15/2023 |
| 47 | 8EBL | 14 | 349 | 2/22/2023 |
| 48 | 8EQ5 | 30 | 305 | 4/26/2023 |
| 49 | 8ESE | 23 | 184 | 5/24/2023 |
| 50 | 8F8Z | 24 | 455 | 1/18/2023 |
| 51 | 8FK3 | 40 | 510 | 3/1/2023 |
| 52 | 8FXQ | 4 | 329 | 2/8/2023 |
| 53 | 8HDJ | 13 | 162 | 5/17/2023 |
| 54 | 8HLO | 13 | 67 | 2/22/2023 |
| 55 | 8I3G | 36 | 184 | 5/31/2023 |
| 56 | 8IA4 | 5 | 101 | 3/22/2023 |
| 57 | 8IA5 | 9 | 90 | 2/22/2023 |
| 58 | 8OEP | 9 | 90 | 5/10/2023 |
| 59 | 8CK5 | 15 | 302 | 4/26/2023 |
| 60 | 8HEP | 10 | 166 | 5/24/2023 |

**Supplementary Table S2.** Details of LOSS values for scoring functions in TB models.

|  | **Mean** | **Median** | **Q1 (25th percentile)** | **Q3 (75th percentile)** | **IQR** | **Minimum** | **Maximum** | **Spread** |
| --- | --- | --- | --- | --- | --- | --- | --- | --- |
| **AFM-Score** | 0.087 | 0.028 | 0.011 | 0.091 | 0.08 | 0.0 | 0.645 | 0.644 |
| **HADDOCK-emscore** | 0.099 | 0.026 | 0.01 | 0.108 | 0.097 | 0.001 | 0.674 | 0.673 |
| **FoldX-Stability** | 0.119 | 0.036 | 0.017 | 0.102 | 0.084 | 0.001 | 0.794 | 0.793 |
| **Vina** | 0.121 | 0.032 | 0.015 | 0.107 | 0.092 | 0.001 | 0.645 | 0.645 |
| **FoldX-Interaction** | 0.126 | 0.042 | 0.015 | 0.169 | 0.154 | 0.001 | 0.771 | 0.771 |
| **HADDOCK-mdscore** | 0.136 | 0.035 | 0.021 | 0.142 | 0.122 | 0.001 | 0.85 | 0.849 |
| **PyRosetta** | 0.138 | 0.036 | 0.015 | 0.139 | 0.123 | 0.001 | 0.876 | 0.875 |
| **Vinardo** | 0.14 | 0.043 | 0.019 | 0.109 | 0.09 | 0.001 | 0.843 | 0.842 |
| **DeepRank-GNN-esm** | 0.162 | 0.047 | 0.016 | 0.198 | 0.182 | 0.001 | 0.864 | 0.864 |
| **GNN_Dove** | 0.164 | 0.049 | 0.022 | 0.226 | 0.204 | 0.0 | 0.846 | 0.846 |

**Supplementary Table S3.** Details of LOSS values for scoring functions in TF models.

|  | **Mean** | **Median** | **Q1 (25th percentile)** | **Q3 (75th percentile)** | **IQR** | **Minimum** | **Maximum** | **Spread** |
| --- | --- | --- | --- | --- | --- | --- | --- | --- |
| **AFM-Score** | 0.101 | 0.047 | 0.015 | 0.121 | 0.107 | 0.0 | 0.67 | 0.67 |
| **FoldX-Stability** | 0.113 | 0.041 | 0.022 | 0.13 | 0.107 | 0.0 | 0.774 | 0.774 |
| **HADDOCK-mdscore** | 0.128 | 0.071 | 0.026 | 0.143 | 0.117 | 0.005 | 0.735 | 0.73 |
| **FoldX-Interaction** | 0.129 | 0.047 | 0.018 | 0.153 | 0.135 | 0.0 | 0.77 | 0.77 |
| **PyRosetta** | 0.133 | 0.041 | 0.021 | 0.177 | 0.156 | 0.0 | 0.844 | 0.844 |
| **HADDOCK-emscore** | 0.134 | 0.051 | 0.017 | 0.158 | 0.141 | 0.003 | 0.728 | 0.726 |
| **Vina** | 0.146 | 0.068 | 0.025 | 0.179 | 0.153 | 0.007 | 0.83 | 0.823 |
| **Vinardo** | 0.147 | 0.081 | 0.032 | 0.199 | 0.168 | 0.002 | 0.843 | 0.842 |
| **DeepRank-GNN-esm** | 0.173 | 0.086 | 0.024 | 0.297 | 0.273 | 0.003 | 0.592 | 0.589 |
| **GNN_Dove** | 0.183 | 0.104 | 0.039 | 0.322 | 0.283 | 0.002 | 0.73 | 0.727 |

**Supplementary Table S4.** Details of Spearman correlation coefficient values for scoring functions in TB models.

|  | **Median** | **Q1 (25th percentile)** | **Q3 (75th percentile)** | **IQR** | **Minimum** | **Maximum** | **Spread** |
| --- | --- | --- | --- | --- | --- | --- | --- |
| **AFM-Score** | 0.133 | -0.312 | 0.497 | 0.809 | -0.807 | 0.915 | 1.722 |
| **GNN_Dove** | 0.123 | -0.357 | 0.313 | 0.67 | -0.811 | 0.887 | 1.698 |
| **DeepRank-GNN-esm** | 0.12 | -0.04 | 0.358 | 0.398 | -0.725 | 0.751 | 1.476 |
| **HADDOCK-emscore** | 0.094 | -0.3 | 0.386 | 0.685 | -0.818 | 0.714 | 1.532 |
| **HADDOCK-mdscore** | 0.088 | -0.126 | 0.241 | 0.367 | -0.543 | 0.76 | 1.303 |
| **PyRosetta** | 0.024 | -0.432 | 0.331 | 0.763 | -0.797 | 0.749 | 1.546 |
| **FoldX-Stability** | 0.001 | -0.283 | 0.302 | 0.585 | -0.78 | 0.625 | 1.405 |
| **Vinardo** | -0.047 | -0.42 | 0.232 | 0.651 | -0.817 | 0.861 | 1.678 |
| **FoldX-Interaction** | -0.071 | -0.388 | 0.24 | 0.629 | -0.762 | 0.719 | 1.481 |
| **Vina** | -0.105 | -0.402 | 0.22 | 0.622 | -0.838 | 0.775 | 1.613 |

**Supplementary Table S5.** Details of Spearman correlation coefficient values for scoring functions in TF models.

|  | **Median** | **Q1 (25th percentile)** | **Q3 (75th percentile)** | **IQR** | **Minimum** | **Maximum** | **Spread** |
| --- | --- | --- | --- | --- | --- | --- | --- |
| **AFM-Score** | 0.182 | -0.247 | 0.521 | 0.768 | -0.847 | 0.913 | 1.76 |
| **GNN_Dove** | 0.123 | -0.076 | 0.315 | 0.391 | -0.759 | 0.701 | 1.46 |
| **HADDOCK-emscore** | 0.094 | -0.198 | 0.333 | 0.531 | -0.714 | 0.786 | 1.5 |
| **FoldX-Stability** | 0.085 | -0.295 | 0.327 | 0.622 | -0.743 | 0.701 | 1.444 |
| **HADDOCK-mdscore** | 0.069 | -0.131 | 0.302 | 0.432 | -0.512 | 0.741 | 1.253 |
| **DeepRank-GNN-esm** | 0.051 | -0.108 | 0.232 | 0.34 | -0.673 | 0.583 | 1.257 |
| **PyRosetta** | 0.041 | -0.255 | 0.368 | 0.623 | -0.768 | 0.706 | 1.474 |
| **FoldX-Interaction** | 0.036 | -0.228 | 0.354 | 0.582 | -0.653 | 0.81 | 1.464 |
| **Vina** | -0.006 | -0.304 | 0.196 | 0.5 | -0.642 | 0.593 | 1.235 |
| **Vinardo** | -0.129 | -0.314 | 0.236 | 0.55 | -0.786 | 0.632 | 1.418 |

**Supplementary Table S6.** Details of common-samples values for three scoring functions in TB models.

|  | **Mean** | **Median** | **Q1 (25th percentile)** | **Q3 (75th percentile)** | **IQR** | **Minimum** | **Maximum** | **Spread** |
| --- | --- | --- | --- | --- | --- | --- | --- | --- |
| **FoldX-HADDOCK** | 0.019 | 0.011 | 0.007 | 0.035 | 0.027 | 0.005 | 0.043 | 0.038 |
| **Common-samples** | 0.075 | 0.019 | 0.007 | 0.124 | 0.117 | 0.004 | 0.248 | 0.243 |
| **AlphaFold-HADDOCK** | 0.1 | 0.028 | 0.009 | 0.177 | 0.168 | 0.0 | 0.529 | 0.529 |
| **AlphaFold** | 0.101 | 0.047 | 0.015 | 0.121 | 0.107 | 0.0 | 0.67 | 0.67 |
| **FoldX-Stability** | 0.113 | 0.041 | 0.022 | 0.13 | 0.107 | 0.0 | 0.774 | 0.774 |
| **HADDOCK-mdscore** | 0.128 | 0.071 | 0.026 | 0.143 | 0.117 | 0.005 | 0.735 | 0.73 |
| **AlphaFold-FoldX** | 0.178 | 0.101 | 0.035 | 0.241 | 0.206 | 0.0 | 0.676 | 0.676 |

**Supplementary Table S7.** Details of common-samples values for three scoring functions in TF models.

|  | **Mean** | **Median** | **Q1 (25th percentile)** | **Q3 (75th percentile)** | **IQR** | **Minimum** | **Maximum** | **Spread** |
| --- | --- | --- | --- | --- | --- | --- | --- | --- |
| **FoldX-HADDOCK** | 0.019 | 0.011 | 0.007 | 0.035 | 0.027 | 0.005 | 0.043 | 0.038 |
| **Common-samples** | 0.075 | 0.019 | 0.007 | 0.124 | 0.117 | 0.004 | 0.248 | 0.243 |
| **AlphaFold** | 0.087 | 0.028 | 0.011 | 0.091 | 0.08 | 0.0 | 0.645 | 0.644 |
| **AlphaFold-HADDOCK** | 0.1 | 0.028 | 0.009 | 0.177 | 0.168 | 0.0 | 0.529 | 0.529 |
| **FoldX-Stability** | 0.119 | 0.036 | 0.017 | 0.102 | 0.084 | 0.001 | 0.794 | 0.793 |
| **HADDOCK-mdscore** | 0.136 | 0.035 | 0.021 | 0.142 | 0.122 | 0.001 | 0.85 | 0.849 |
| **AlphaFold-FoldX** | 0.178 | 0.101 | 0.035 | 0.241 | 0.206 | 0.0 | 0.676 | 0.676 |

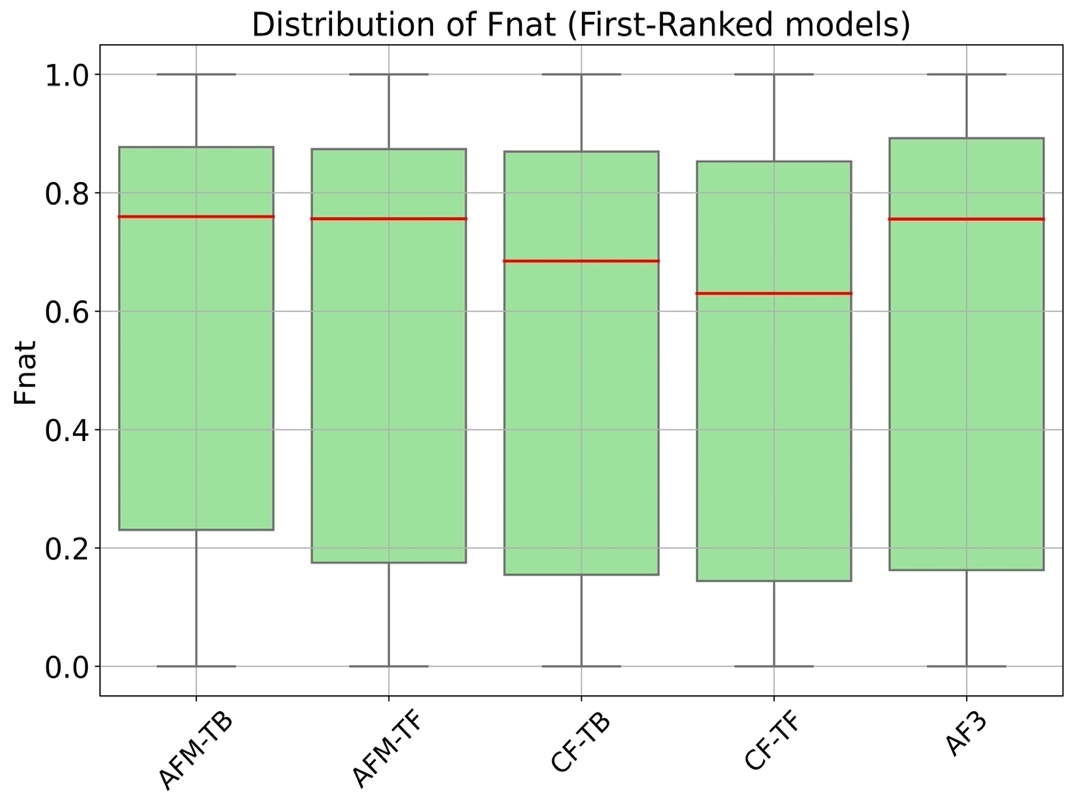

**Fig. S1. Fnat (Fraction of Native Contacts) on the first-ranked models.** Fnat measures the preservation of native contacts in predicted protein structures, with higher values indicating better accuracy. AFM-TB had the highest median Fnat (0.75) and the narrowest IQR (0.64), indicating high-quality and consistent predictions. AFM-TF (median: 0.75, IQR: 0.69) and AF3 (median: 0.75, IQR: 0.72) also performed well but with more variability. CF-TB (median: 0.68, IQR: 0.71) and CF-TF (median: 0.63, IQR: 0.70) had lower medians, indicating less accuracy.

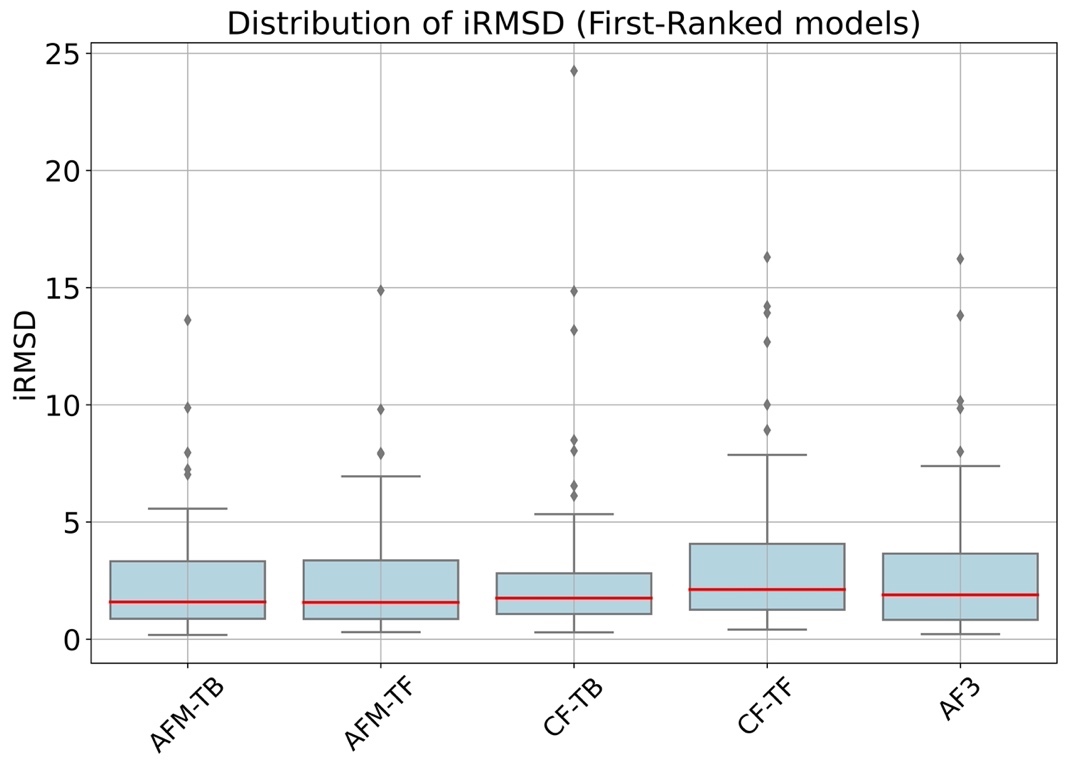

**Fig. S2. iRMSD (Interface Root Mean Square Deviation) on the first-ranked models.** iRMSD measures the deviation between predicted and reference structures, with lower values indicating better accuracy. AFM-TF had the lowest median iRMSD (1.57) with moderate variability (IQR: 2.50), indicating good accuracy. AFM-TB had a similar median (1.58) and a slightly narrower IQR (2.45), suggesting consistent predictions. CF-TB (median: 1.75, IQR: 1.73) and AF3 (median: 1.8945, IQR: 2.82) showed higher medians with varying consistency. CF-TF had the highest median (2.12) and the widest IQR (2.81), indicating less accurate predictions. Therefore, AFM-TF and AFM-TB are the best tools based on low and consistent iRMSD values, while CF-TF requires improvement.

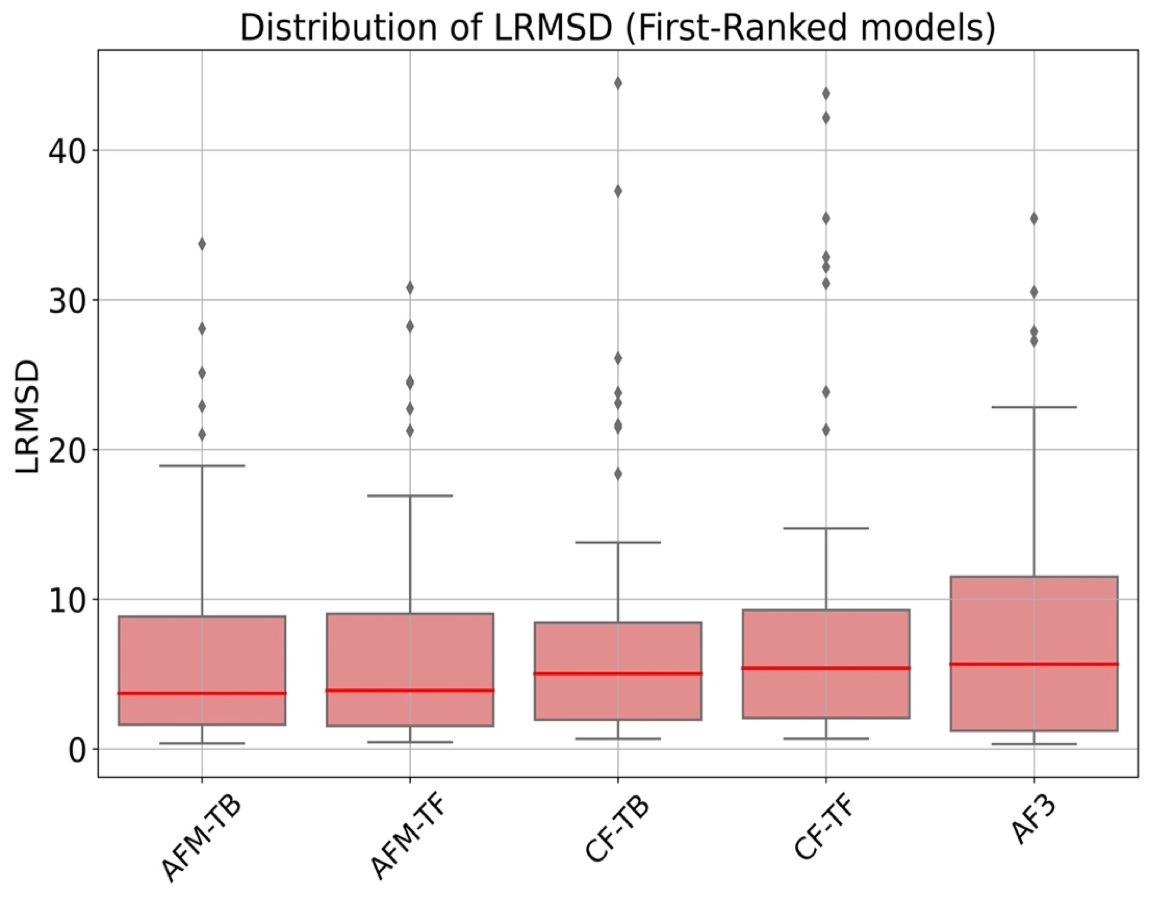

**Fig. S3. LRMSD (Ligand Root Mean Square Deviation) on the first-ranked models.** LRMSD measures the deviation of the ligand's backbone atoms between predicted and reference structures, with lower values indicating better accuracy. AFM-TB had a median LRMSD of 3.70 with an IQR of 7.22, showing moderate accuracy and variability. AFM-TF had a slightly higher median (3.90) and IQR (7.49), indicating similar performance. CF-TB (median: 5.03, IQR: 6.47) and CF-TF (median: 5.39, IQR: 7.22) showed higher medians, suggesting lower accuracy. AF3 had the highest median LRMSD (5.65) and the widest IQR (10.29), indicating the least accurate and most variable predictions. Therefore, AFM-TB and AFM-TF are the best tools based on their lower and relatively consistent LRMSD values, while AF3 requires the most improvement.

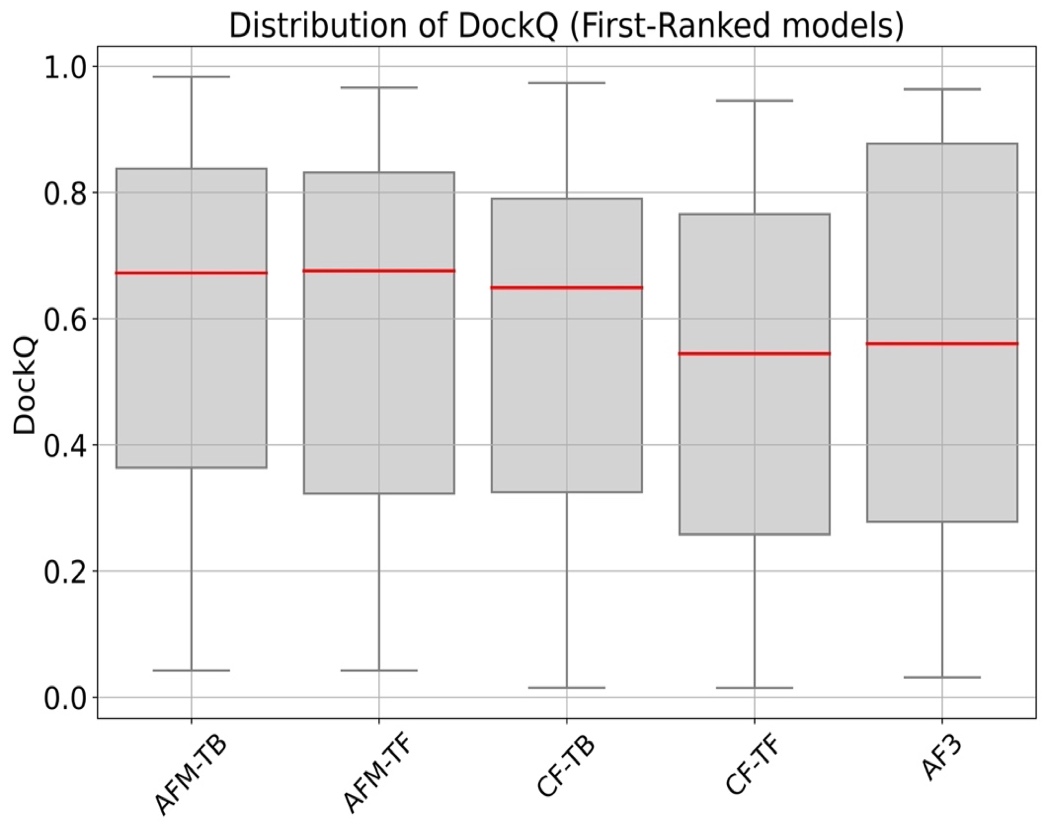

**Fig. S4. DockQ on the first-ranked models.** DockQ is a composite score measuring the quality of predicted protein-protein interactions, with higher values indicating better predictions. AFM-TF had the highest median DockQ score (0.67) with an IQR of 0.50, indicating good accuracy and moderate variability. AFM-TB had a similar median (0.67) and a slightly narrower IQR (0.47), suggesting consistent performance. CF-TB (median: 0.64, IQR: 0.46) also performed well but with slightly lower accuracy. CF-TF (median: 0.54, IQR: 0.50) and AF3 (median: 0.56, IQR: 0.59) showed lower medians and wider variability, indicating less accurate predictions. Therefore, AFM-TF and AFM-TB are the best tools based on their high and consistent DockQ values, while CF-TF and AF3 require improvement.

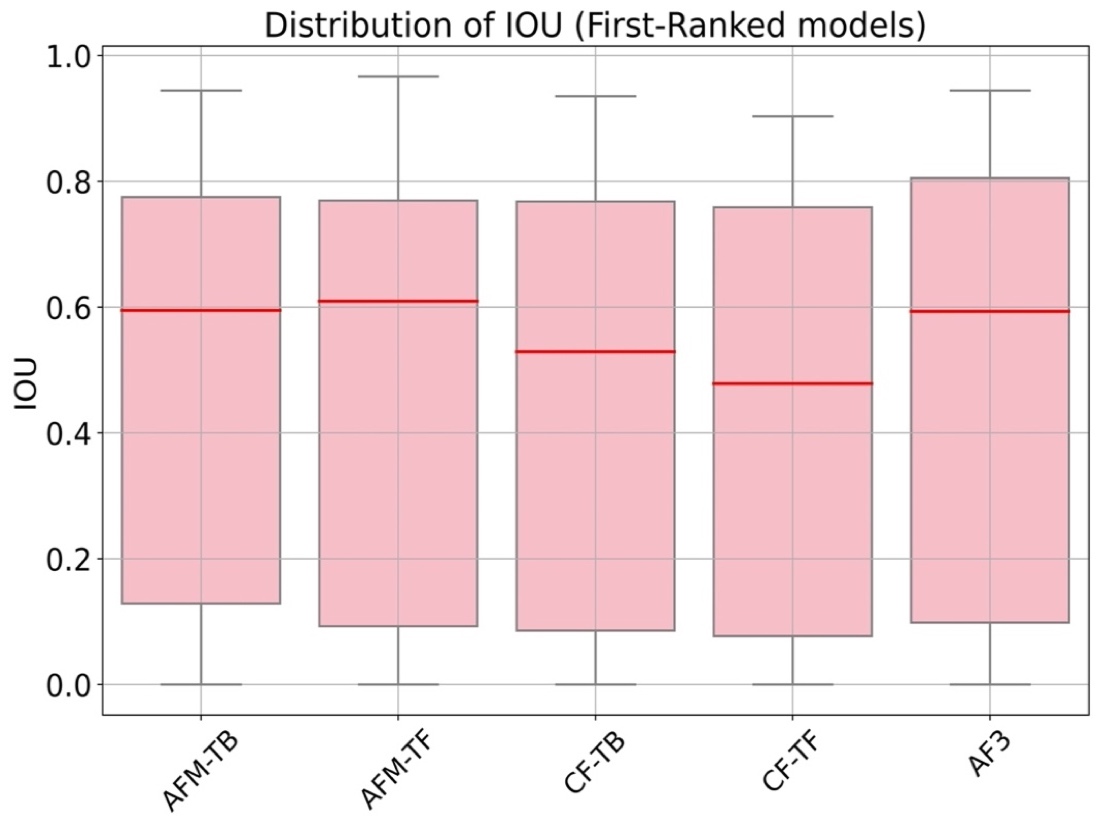

**Fig. S5. IOU (Intersection Over Union) on the first-ranked models.** IOU measures the overlap between predicted and reference protein-protein interaction interfaces, with higher values indicating better predictions. AFM-TF had the highest median IOU (0.60) and an IQR of 0.67, indicating good accuracy with moderate variability. AFM-TB followed closely with a median of 0.59 and a narrower IQR of 0.64, suggesting consistent performance. AF3 also performed well with a median of 0.59 and the widest IQR of 0.70, indicating more variability. CF-TB (median: 0.52, IQR: 0.68) and CF-TF (median: 0.47, IQR: 0.68) had lower medians, indicating less accurate predictions. Therefore, AFM-TF and AFM-TB are the best tools based on their higher and consistent IOU values, while CF-TF requires the most improvement.

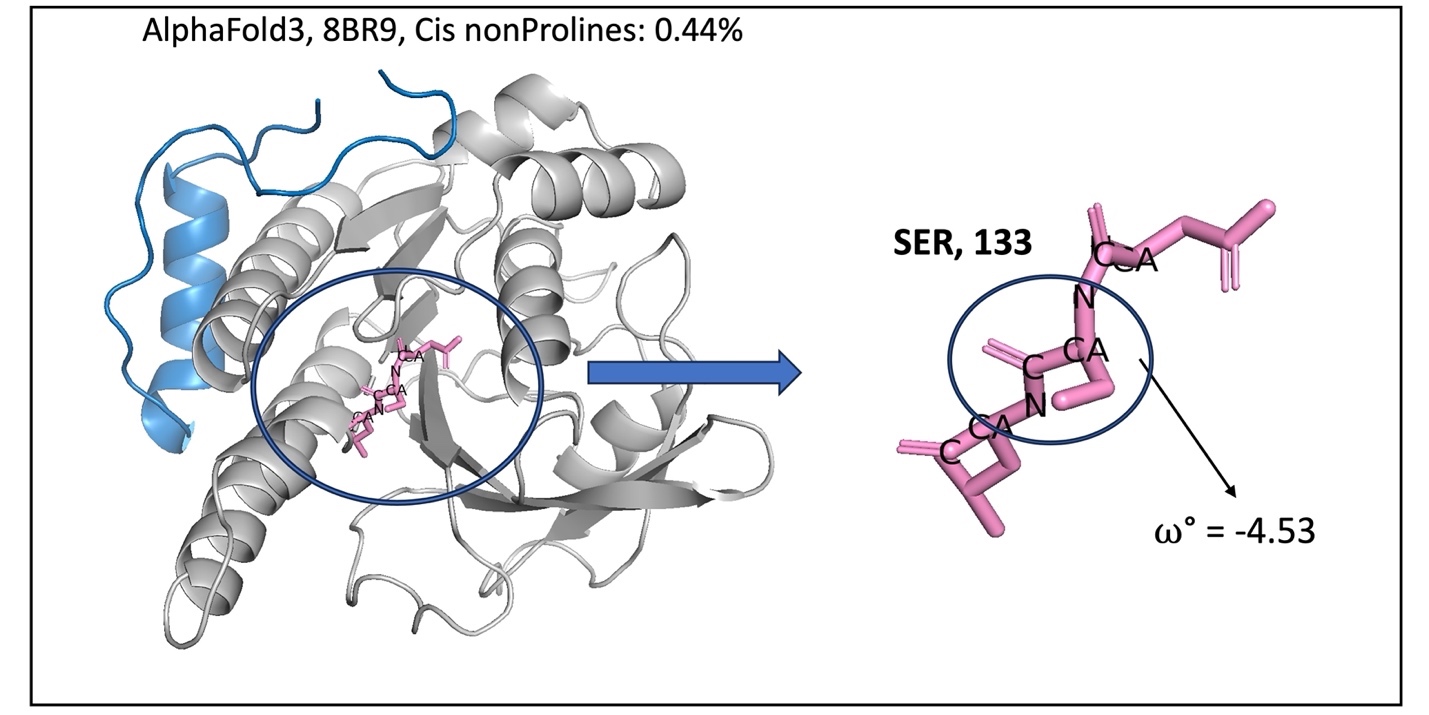

**Fig. S6. Sample (8BR9) of cis non-proline peptide predicted by AF3.** The observed cis non-proline peptide occurred in the protein part, with no cis non-proline peptide identified in the peptide part.

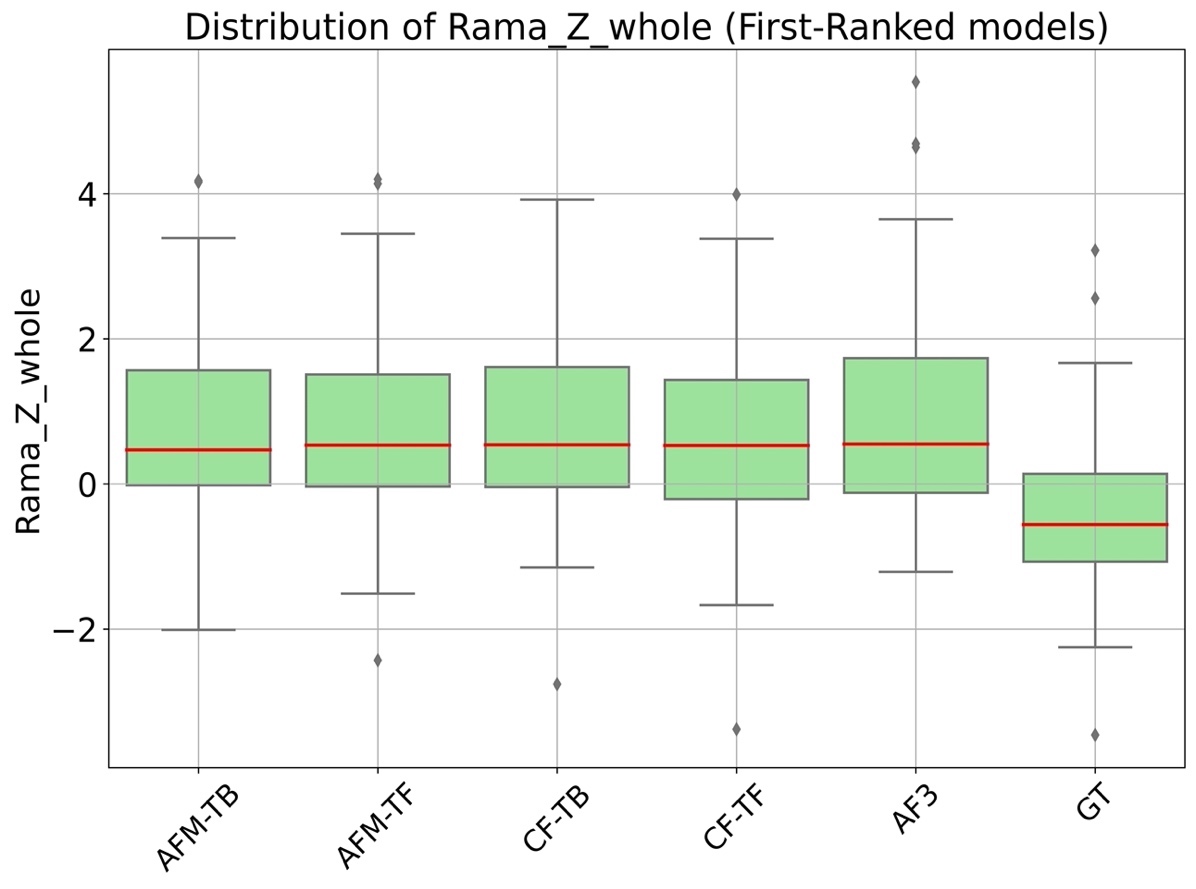

**Fig. S7. Distribution of Rama_Z_whole Scores across different protein modeling methods.** Rama_Z_Score between -2, 2 and close to zero indicate accurate, standard geometries, while scores outside the range of -2 to 2, particularly beyond an absolute value of 3, suggest improbable structures. The Z-scores (Rama-Z) for predicted structures show positive median values ranging from 0.47 to 0.55, indicating they are within the acceptable range (|Rama-Z| < 2). In contrast, the GT median is -0.55, also within the good range, but with a significantly different distribution from the predicted models. The predicted models exhibit greater variability, with IQRs ranging from 1.54 to 1.85, compared to the GT, IQR of 1.17, indicating less consistency. Significant outliers are present in the predicted models, highlighting areas for improvement.

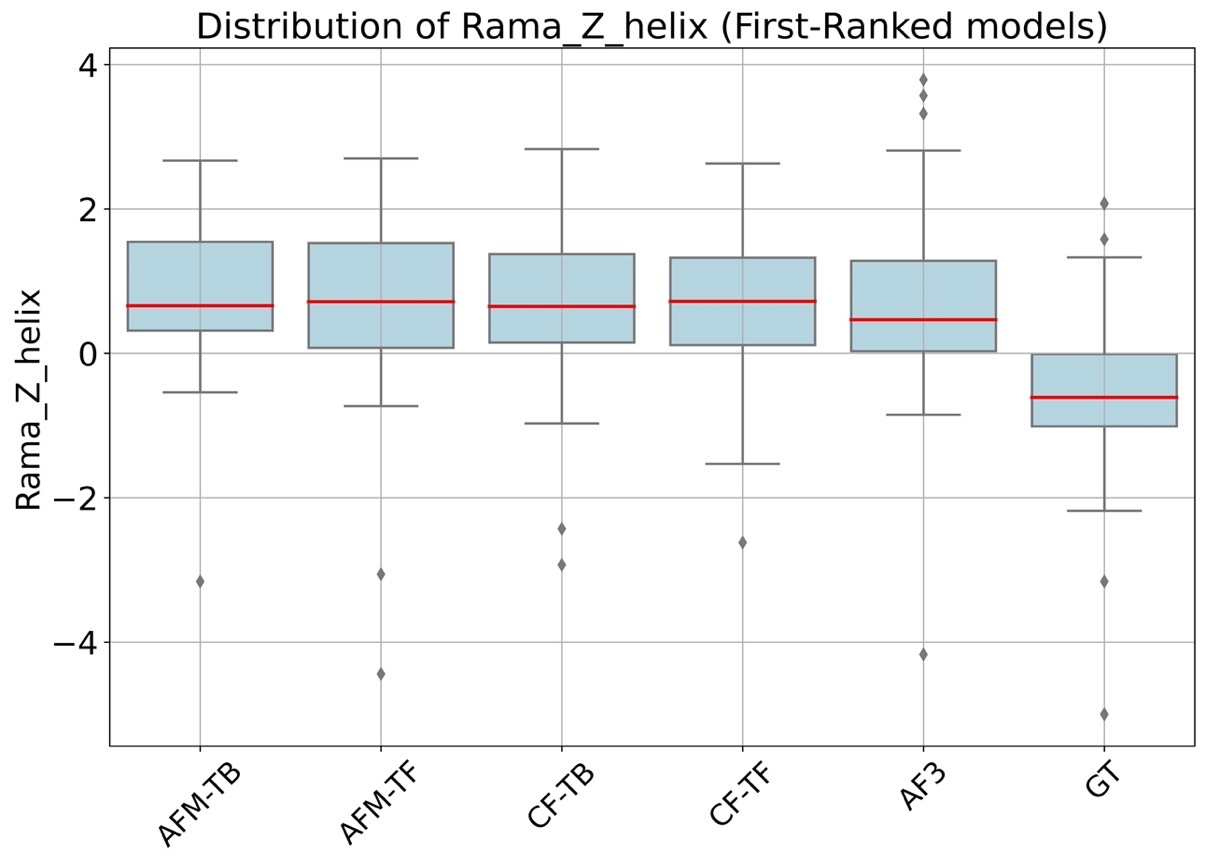

**Fig. S8. Distribution of Rama_Z_helix Scores.** The analysis of Ramachandran plot Z-scores (Rama-Z) for helix regions reveals that predicted models have positive median values ranging from 0.46 to 0.72, indicating good dihedral angles, whereas the GT median is -0.60, also within the acceptable range. Predicted models display higher variability, with IQRs ranging from 1.21 to 1.45, compared to the GT IQR of 1.03. Significant outliers are present in both minimum (-4.44 to -2.93) and maximum (2.63 to 3.79) values for the predicted models. The GT also shows notable outliers, with minimum and maximum values of -5.0 and 2.08, respectively, and a spread of 7.08, comparable to AFM-TF (7.14) and AF3 (7.96). In some samples, MolProbity could not find the value for Rama_Z_helix parameters in both the TB and TF models such as: 8ebl, 8hlo, 7ui8, 7zax.

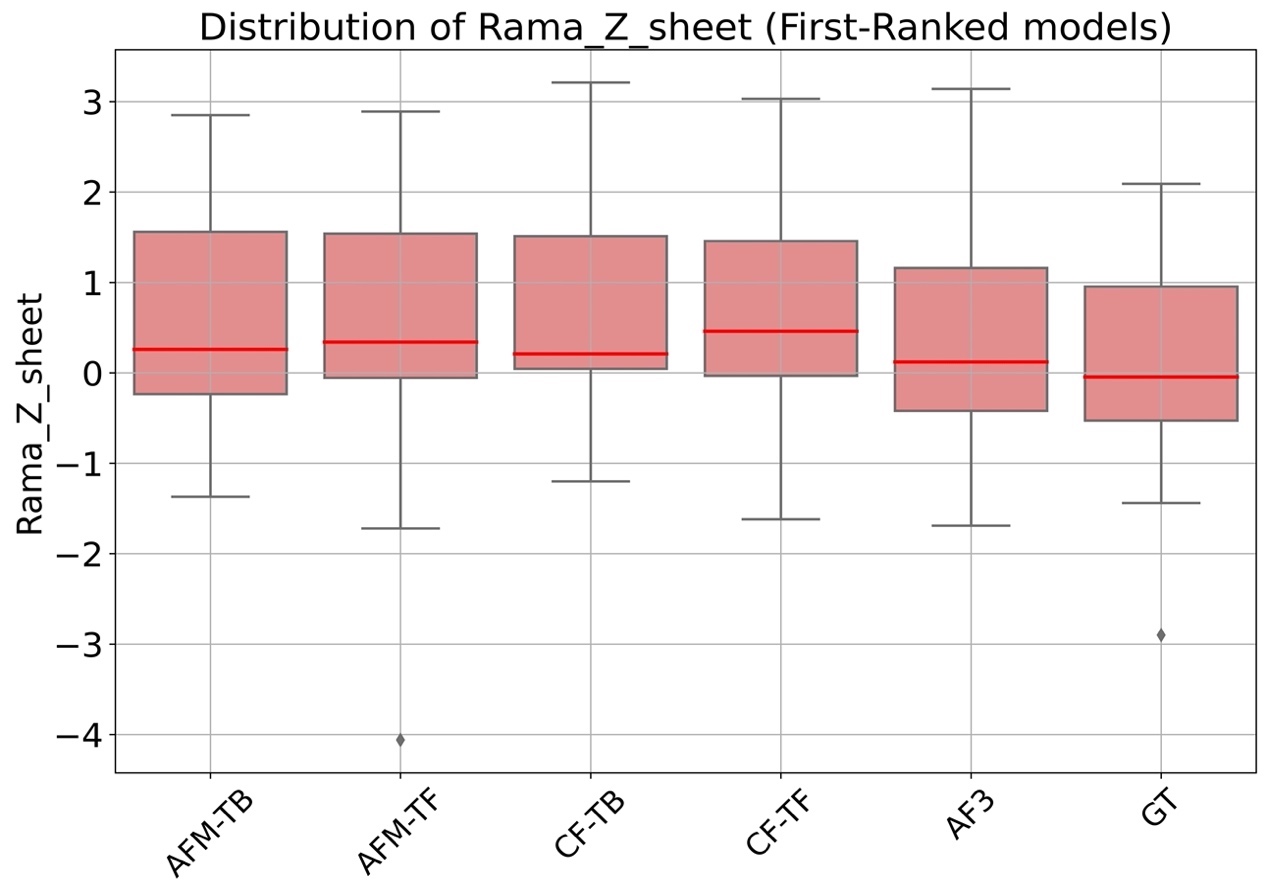

**Fig. S9. Distribution of Rama_Z_sheet Scores.** The analysis of Ramachandran plot Z-scores (Rama-Z) for sheet regions reveals that predicted models have positive median values (0.12 to 0.46), indicating generally good dihedral angles. In contrast, the GT data has a slightly negative median (-0.09), also within the good range. Predicted models show higher variability, with interquartile ranges (IQR) from 1.46 to 1.79, compared to the GT IQR of 1.50, indicating greater consistency in the GT data. Minimum values for predicted models range from -4.06 to -1.20, and maximum values range from 2.85 to 3.21, whereas the GT values range from -2.90 to 2.09. Spreads for predicted models vary from 4.22 (AFM-TB) to 6.95 (AFM-TF), with the GT showing a spread of 4.99. This comparison highlights that while predicted models are accurate, they exhibit greater variability and more extreme outliers than the GT data, suggesting room for refinement to better align with native sheet conformations. In some samples, MolProbity could not find the value for Rama_Z_sheet parameters in both the TB and TF models (7prx, 7sqa, 7t09, 7udk, 7udl, 7ue2, 7uw2, 7v1a, 7wqq, 7xfg, 7y8f, 7z5l, 7z7c, 7zw4, 8a68, 8ahs, 8czk, 8dgm, 8fk3, 8i3g).

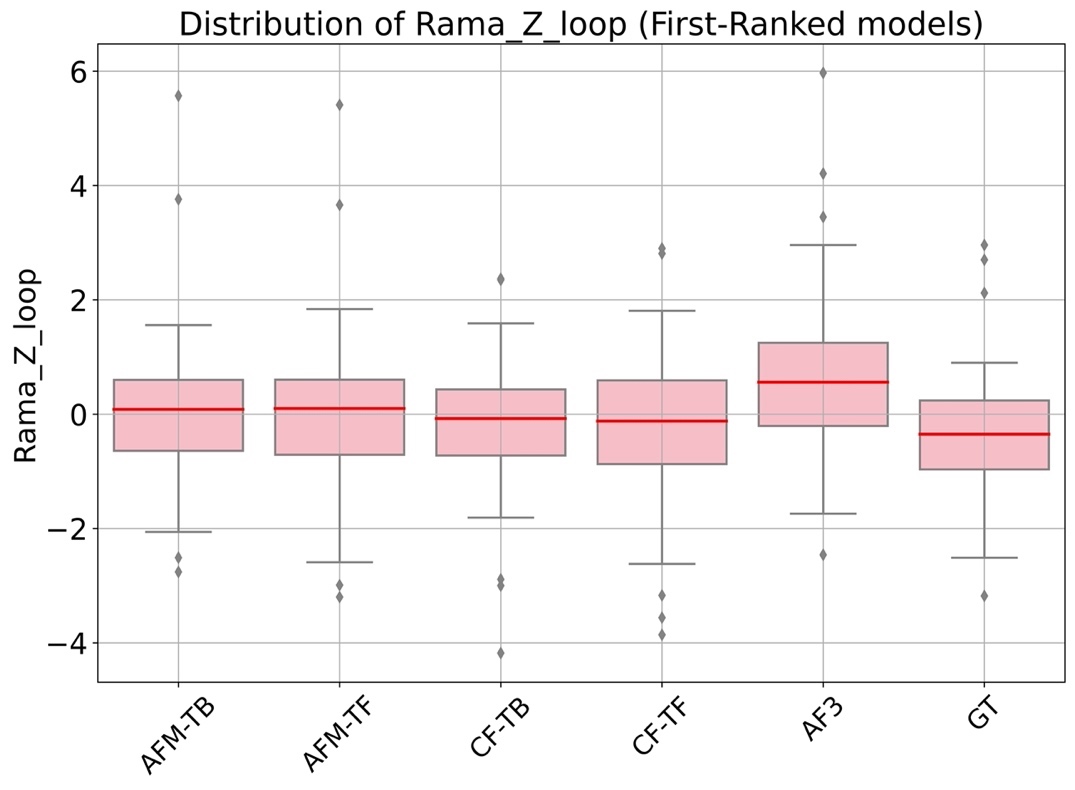

**Fig. S10. Distribution of Rama_Z_loop Scores.** The analysis of Ramachandran plot Z-scores (Rama-Z) for loop regions shows that predicted models have median values close to zero, indicating generally accurate dihedral angles. Specifically, the median values are 0.08 for AFM-TB, 0.1 for AFM-TF, -0.075 for CF-TB, -0.12 for CF-TF, and 0.56 for AF3, compared to the GT median of -0.24. Predicted models exhibit higher variability, with IQRs ranging from 1.15 to 1.46 and wider spreads from 6.55 to 8.61, compared to the GT IQR of 1.22 and spread of 6.14. Minimum values for predicted models range from -4.18 (CF-TB) to -2.46 (AF3), and maximum values range from 2.37 (CF-TB) to 5.97 (AF3), whereas the GT values range from -3.18 to 2.96.

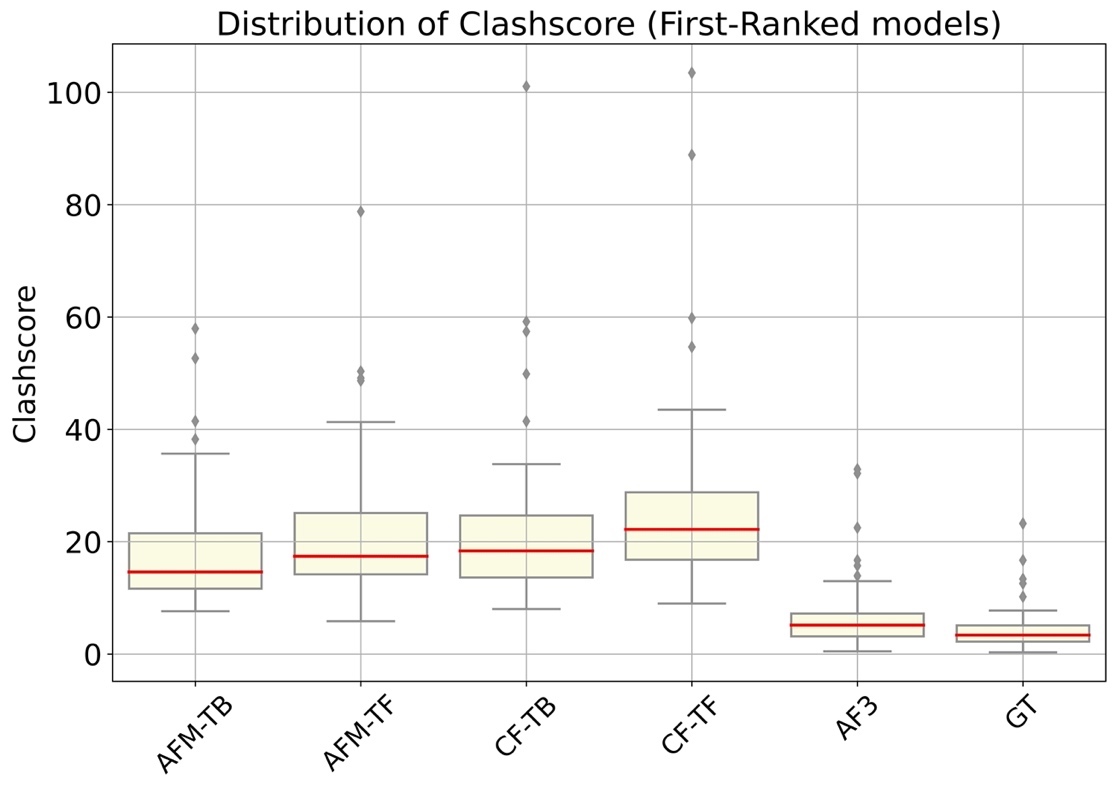

**Fig. S11. Distribution of Clashscore values across different prediction methods.** The analysis of Clashscore, where a lower score indicates better structural quality, reveals significant differences between predicted structures (AFM-TB, AFM-TF, CF-TB, CF-TF, AF3) and GT. GT exhibits the highest structural quality with the lowest median Clashscore of 3.39 and the smallest IQR of 3.09, indicating consistent and high-quality structures. Among predicted models, AF3 stands out with a relatively low median Clashscore of 5.14 and an IQR of 4.06, though still higher than GT. AFM-TB and AFM-TF have higher medians of 14.62 and 17.42, respectively, with wider spreads, indicating more variability and a higher occurrence of steric clashes. CF-TB and CF-TF show the highest medians of 18.36 and 22.18, respectively, along with the largest spreads (93.04 and 94.47), suggesting significant room for improvement. GT’s minimum and maximum values (0.31 to 13.35) reflect fewer extreme outliers than predicted models, with AF3 being the closest (0.49 to 32.89). Overall, while predicted models demonstrate reasonable accuracy, they require refinement to reduce steric clashes and achieve structural quality closer to that of the GT.

**a.**

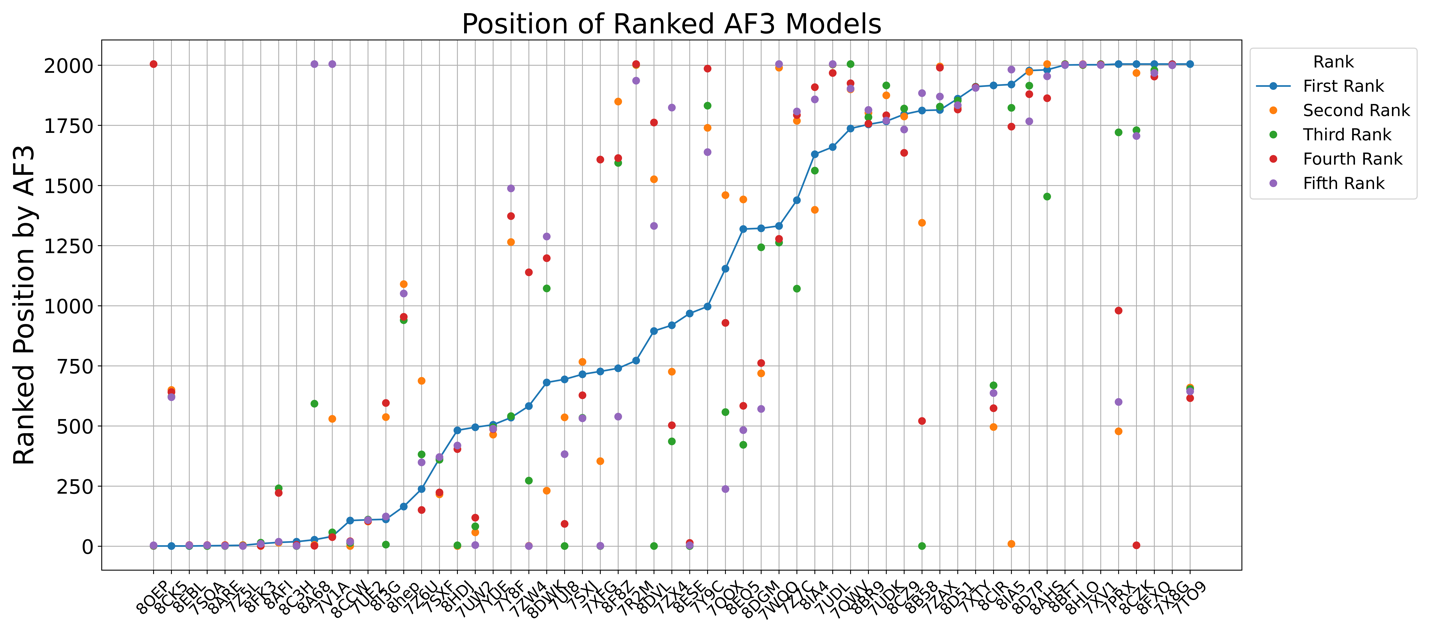

**b.**

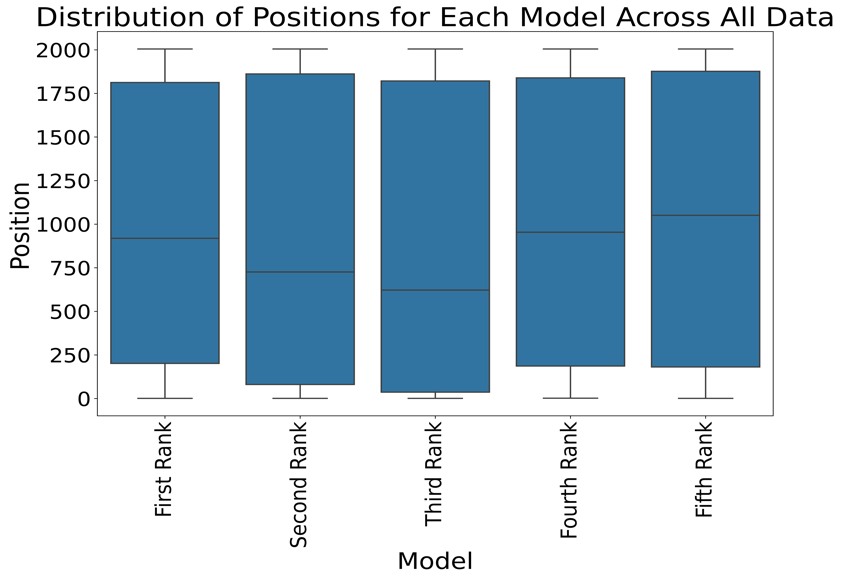

**Fig. S12. Analyzing the ranking positions of AF3.** a. Line graph showing rankings for first-ranked AF3 models within a pool of 2005, including 1000 TB and 1000 TF models from AFM, and dots illustrating the ranking of other AF3’s ranked models. b. Box plot illustrated the ranking distributions, highlighting the third AF3 model's superior performance with a lower median than other ranks. AFM generally predicts nearer-native structures more effectively in the combined pool. The number of top 10 ranking positions for predicted structures with AF3 is calculated for each model as follows: First Rank - 6, Second Rank - 12, Third Rank - 13, Fourth Rank - 8, Fifth Rank - 10. These numbers indicate how often each model appears in the top ten ranking positions among all samples in the pool. This suggests that AFM structures mostly appear in the top ten ranking positions within the pool. Additionally, the Third and Second-ranked models of AF3 perform better than its First rank positions based on these numbers.

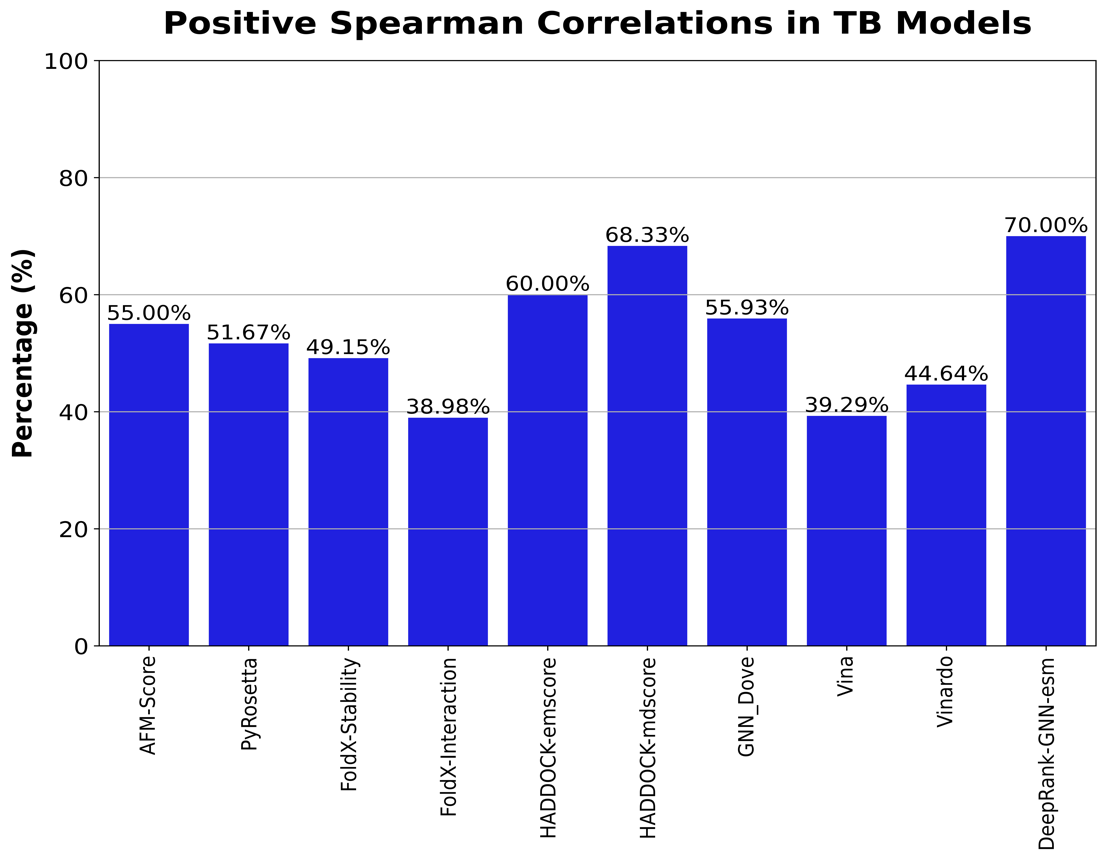

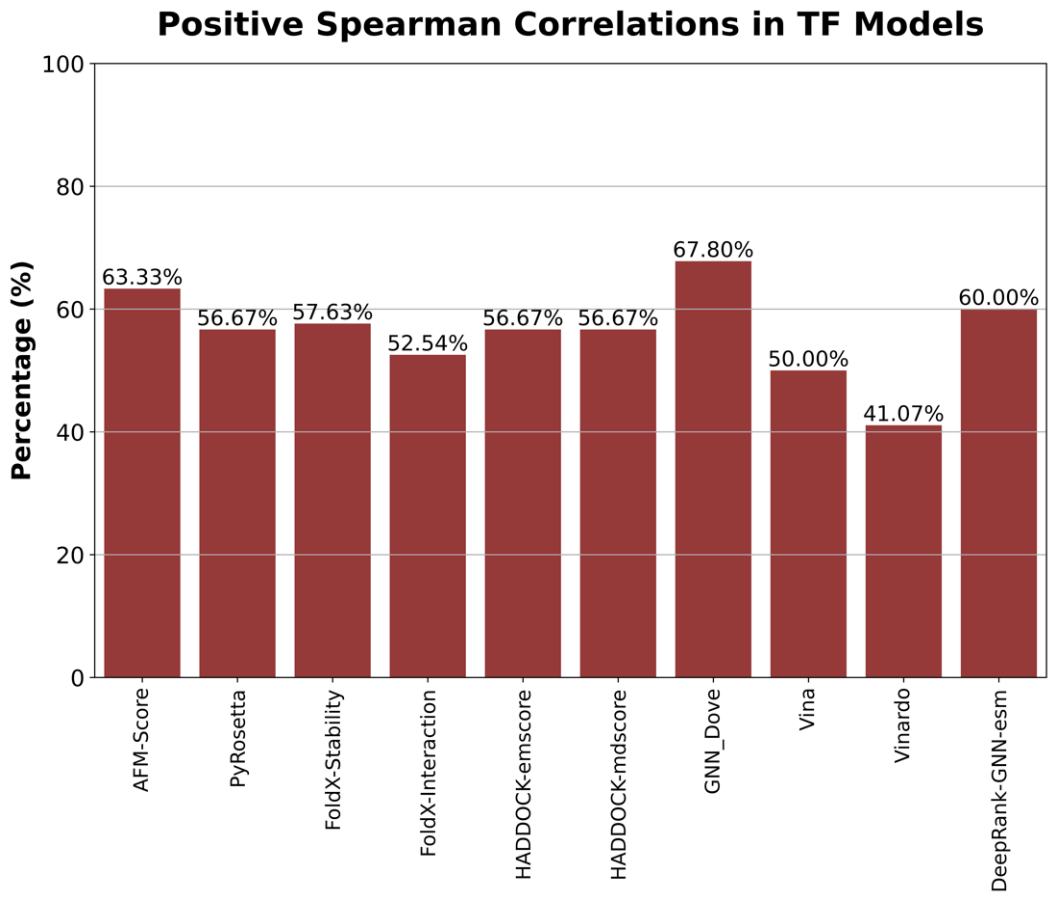

**Fig. S13. Positive Spearman correlation coefficient.** It evaluates scoring functions' capability to rank TB models against the DockQ standard. DeepRank-GNN-esm, with a 70% alignment, ranks as the most consistent with DockQ, showcasing superior accuracy in protein structure quality predictions. Following closely, HADDOCK-emscore and HADDOCK-mdscore rank the structures at 68.33% and 60%, respectively. Other functions like GNN_Dove marginally outperform AFM-Score, recording values of 55.93%. In contrast, PyRosetta and FoldX-Stability show slightly lower performance than AFM-Score at 51.67% and 49.15%, respectively, while FoldX-Interaction, at 38.98%, demonstrates the lowest correlation in rankings. The TF plot examines the ability of scoring functions to rank TF models, highlighting GNN_Dove's top correlation with DockQ at 67.8%. DeepRank-GNN-esm with 60% closely follows, matching AFM-Score’s performance at 63.33%. Other scoring functions fall short of AFM-Score’s effectiveness, with Vinardo showing the lowest ranking capability in TF models.
